# Stromal and Endothelial Transcriptional Changes during Progression from MGUS to Myeloma and after Treatment Response

**DOI:** 10.1101/2024.04.24.589777

**Authors:** Itziar Cenzano, Miguel Cócera, Marta Larrayoz, Lorea Campos-Dopazo, Sonia Sanz, Azari Bantan, Amaia Vilas-Zornoza, Patxi San-Martin, Paula Aguirre-Ruiz, Diego Alignani, Aitziber Lopez, Ignacio Sancho González, Javier Ruiz, Purificacion Ripalda-Cemborain, Marta Abengozar-Muela, Emma Muiños-López, Vincenzo Lagani, Jesper Tegner, Mikel Hernáez, Xabier Agirre, Benjamin Ebert, Bruno Paiva, Paula Rodriguez-Otero, Luis-Esteban Tamariz-Amador, Jesús San-Miguel, Borja Saez, José A. Martinez-Climent, Isabel A. Calvo, David Gomez-Cabrero, Felipe Prosper

## Abstract

The role of the non-immune bone marrow microenvironment (BME) in the transition from monoclonal gammopathy of undetermined significance (MGUS) into clinically active multiple myeloma (MM) remains incompletely defined. To address this, we transcriptionally profiled endothelial cells (EC), mesenchymal stem cells (MSC) and MM cells at single-cell resolution from two genetically engineered mouse models (BI_cγ1_ and MI_cγ1_) that recapitulate MGUS to MM progression. Our analysis revealed distinct transcriptional trajectories in EC and MSC, uncovering stage-specific BME–PC interactions shaping disease progression. EC acquired a stress phenotype during MGUS transitioning to angiogenesis in MM, while MSC exhibited early impaired differentiation capacity during MGUS that persisted in MM. Notably, an interferon (IFN)-associated MM signature was detected in EC and MSC from the BI_cγ1_ model but was absent in the more aggressive MI_cγ1_ model. Treatment with bortezomib, lenalidomide, and dexamethasone remodeled the BME by suppressing MM-IFN signaling, promoting an adaptive response in EC, and restoring osteogenic potential in MSC— shifting the niche toward a less tumor-permissive state. Importantly, the MM-IFN signature was validated in patients across the MGUS-to-MM spectrum, supporting the translational relevance of our findings. Together, these data define novel dynamic and targetable alterations in the non-immune BME during myeloma progression.

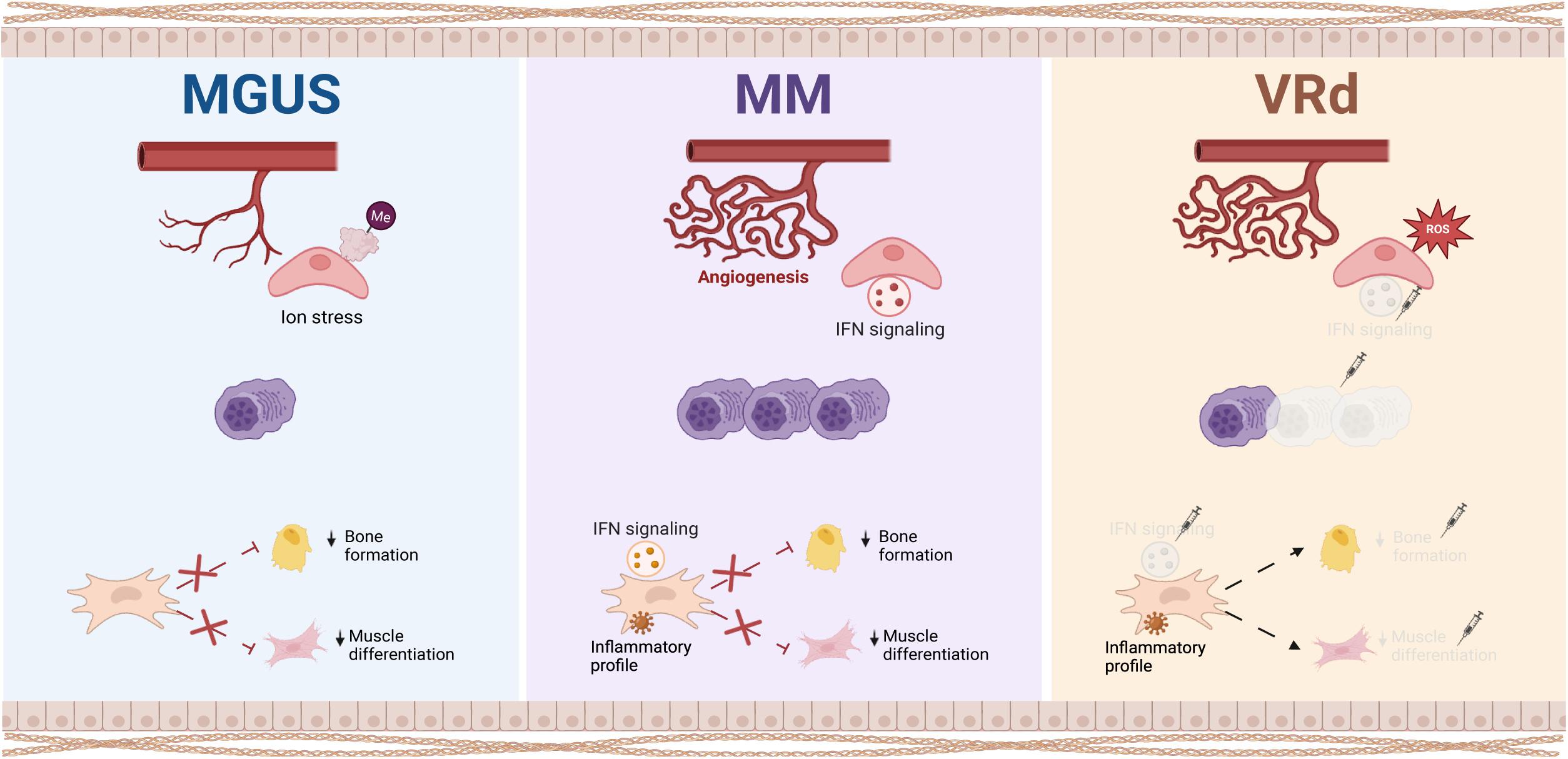

## INTRODUCTION

Multiple myeloma (MM) is a B-cell malignancy characterized by the clonal expansion of plasma cells (PC) in the bone marrow (BM), preceded by asymptomatic precursor stages: monoclonal gammopathy of undetermined significance (MGUS) and smoldering multiple myeloma (SMM)^1^. Understanding the molecular mechanisms driving disease progression is crucial for developing therapies to prevent symptomatic MM.

The BM microenvironment (BME) comprises a complex network of hematopoietic and non-hematopoietic cells, extracellular matrix components, and soluble factors. This microenvironment creates an immunosuppressive and proinflammatory milieu that supports malignant PC growth and survival^2^. Therefore, the intricate interactions between myeloma cells and their surrounding microenvironment are crucial for developing effective therapies that can disrupt supportive signals and overcome drug resistance. While most studies have focused on the immune component^3–6^, increasing evidence suggests that endothelial cells (EC) and mesenchymal stem cells (MSC) foster myeloma PC^7–9^. Furthermore, an enhanced inflammatory signature has been identified in MSC of MM patients^10–12^. However, the role of the non-hematopoietic BME to MGUS-to-MM progression remains poorly understood.

In this study, we took advantage of our recently described mouse models of MM that recapitulate the clinical, genetic, molecular, and immunological characteristics of MM patients^13^. These models include a precursor stage MGUS-like that progresses into symptomatic MM. Using single-cell RNA sequencing (scRNA-seq) data of Fluorescent-Activated Cell Sorted (FACS) non-hematopoietic BME cells and PC, we analyzed the transcriptional changes during disease transition and in response to current therapy. Our results identified distinct transcriptional dynamics between EC and MSC at the different stages of the disease. EC exhibited an altered vascular profile at MGUS, followed by the upregulation of angiogenesis pathways during MM. In contrast, MSC exhibited early downregulation of pathways associated with adhesion, osteogenic, and myogenic differentiation at the MGUS stage. Interestingly, we identified an interferon (IFN)-related myeloma signature in both EC and MSC that was reversed after treatment with bortezomib (Velcade®), lenalidomide (Revlimid®), and dexamethasone (VRd). Changes were validated in MSC from patients with MGUS, SMM, and MM, highlighting their potential role in disease progression.

## METHODS

All detailed information is provided in supplemental methods.

### Mice

We used two genetically engineered MM mouse models, BI_cγ1_ and MI_cγ1_, that recapitulate the common features observed in human MM. YFP_cγ1_ mice were used as controls^13^ (**Supplementary Table 1**). For each scRNA-seq experiment at MGUS and MM-like stages, we used a pool of BM cells from 4-6 mice. In BI_cγ1_ mice with a humanized cereblon (CRBN)^14^, VRd therapy was administered by oral gavage for 20 consecutive weeks. The study included mice of both sexes and was approved by the Ethical Committee of Animal Experimentation of the University of Navarra and the Health Department of the Navarra Government.

Murine BM non-hematopoietic cells were isolated as previously described^15^ and sorted with GFP-positive (GFP^+^) PC using BD FACSAria II for subsequent scRNA-seq.

### Human

BM aspirates from individuals of both sexes with newly diagnosed MGUS (n = 5), SMM (n =2), or MM (n =10), and bone-chip BM samples from healthy donors (HD) (50-80 years of age) HD (n=8) undergoing orthopedic surgery were collected for bulk RNA-sequencing (bulk RNA-seq). Immunofluorescence (IF) staining was performed on human BM biopsies from MGUS (n=4) and MM (n=7) patients, as well as on bone chips from HD (n=2). Written informed consent was obtained according to the Declaration of Helsinki and the Institutional Review Board of the University of Navarra. The clinical characteristics of all individuals are described in **Supplementary Table 2**.

Samples were processed and BM cells were cryopreserved with FBS and 10% dimethyl sulfoxide (Sigma) in liquid nitrogen until use. Frozen samples were thawed, and MSC were prospectively isolated and stored at - 80°C for bulk RNA-seq.

### Single-cell RNA and Bulk RNA sequencing

Details on library preparation, sequencing, and additional information, including differential expression analysis (DEA), gene ontology (GO) over-representation analysis (ORA), gene set enrichment analysis (GSEA), cell-cell communication, and gene regulatory network (GRN) analysis, are described in supplemental methods.

## RESULTS

### Single-cell transcriptomic profiling of BM murine cells in control, MGUS, and MM

Using the BI_cγ1_ myeloma model^13^, we aimed to characterize transcriptional changes in BM stromal and EC during transition from normal and precursor conditions to MM. This model, which carries BCL2 and IKK2^NF-κB^ genetic lesions triggered in germinal center B cells by a cre-cγ1 allele, develops signs of MGUS-like around 6 months of age that progressively evolves into MM within 4 to 6 months, overcoming limitations of previous models. Considering that BI_cγ1_ mice received T-cell-driven immunization, we included similarly immunized YFP_cγ1_ mice, carrying a yellow-fluorescence reporter activated from germinal-center B cells, as controls (**Fig. 1A**). We performed scRNA-seq on FACS-sorted non-hematopoietic BME cells (Lin^-^, TER119^-^, CD45^-^, VDO^+^, and 7AAD^-^) and PC (VDO^+^, 7AAD^-^ and GFP^+^), yielding 12 437 cells (control=2 416, MGUS=4 169, MM=5 852) (**Supplementary Fig. 1**). Using canonical markers, we identified 17 clusters comprising EC, MSC, PC, and other immune cells from the BME (**Fig. 1B**, **Supplementary Fig. 2A,B**). As a result, eight clusters were labeled as PC (clusters PC1-8; n=7 924; markers: *Sdc1*, *Xbp1, Tnfrsf17,* and *Slamf7*); two clusters as erythroblasts (Erythro1-2; n=457; *Hba-a1*, *Hba-a2, Car1,* and *Car2*); one cluster as neutrophils (n=176; *S100a8*, *S100a9, Fcer1g,* and *Lyz2*); three clusters as EC (EC1-3; n= 3 288; *Cdh5*, *Cav1*, *Ly6c1*, and *Kdr*); and two clusters as MSC (MSC1-2; n= 592; mesenchymal and osteolineage-primed markers *Cxcl12*, *Pdgfra*, *Lepr*, and *Col1a2*) (**Fig. 1C, Supplementary Fig. 2C-F**).

**Fig. 1.**
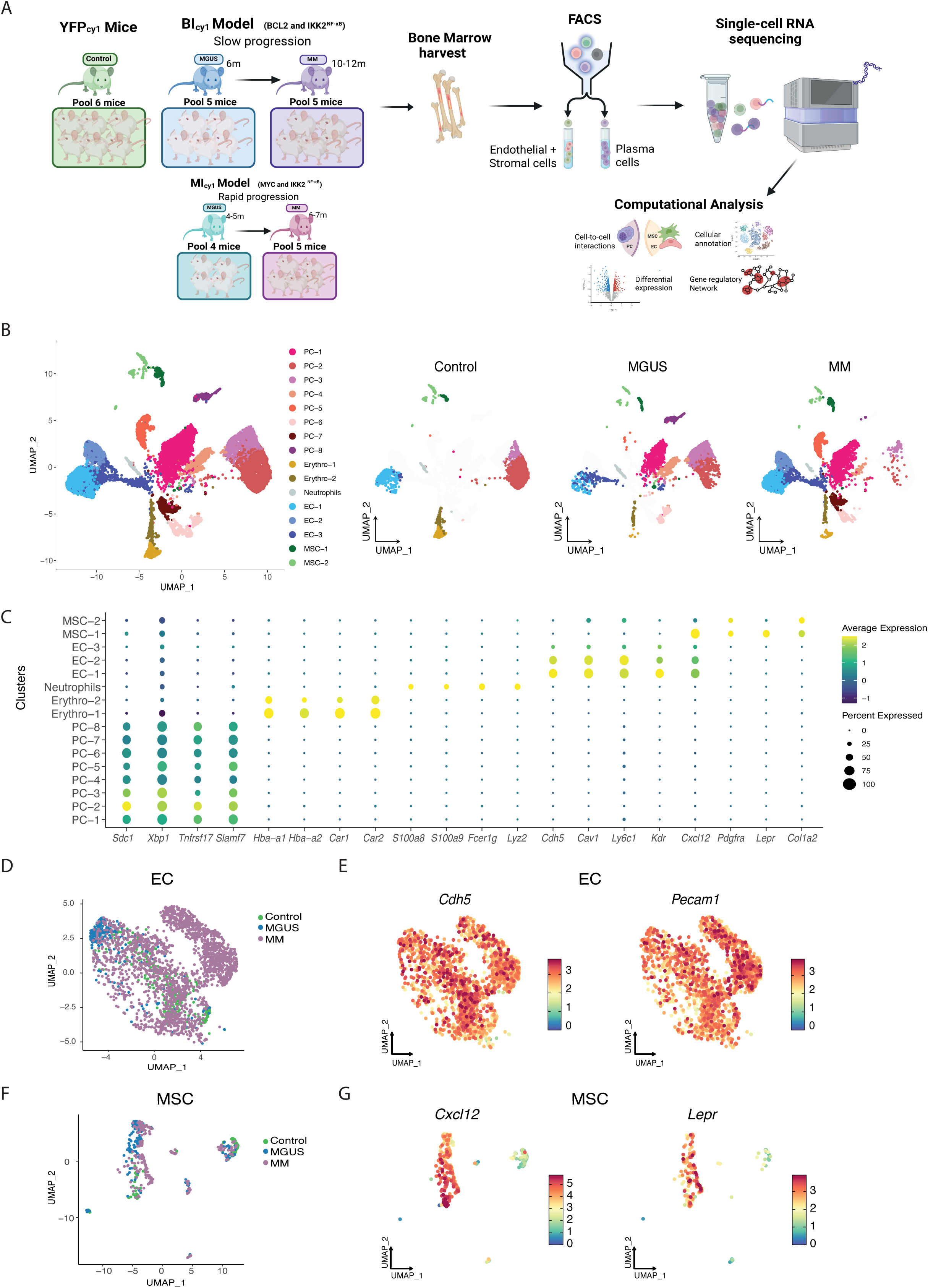
Single-cell transcriptomic profiling of BM murine cells in control, MGUS, and MM. (A) Experiment overview. Schematic view of experimental workflow for the isolation and scRNA-seq analysis of PC, MSC and EC from control (YFP_cγ1_) (pool of 6 mice), MGUS BI_cγ1_ (pool of 5 mice), MM BI_cγ1_ (pool of 5 mice), MGUS MI_cγ1_ (pool of 4 mice) and MM MI_cγ1_ (pool of 5 mice) murine models. (B) Left panel: UMAP representation of murine BM microenvironment control, MGUS, and MM cells colored by cell clustering after merging samples. Right panels: Original UMAP split into control, MGUS, and MM datasets. (C) Dot plot of canonical markers used to define PC, EC, MSC, erythroblasts, and neutrophils. The dot size reflects the percentage of cells within the cluster expressing each gene, and the color represents the average expression level. (D) UMAP projection showing the distribution of EC from control (n=315 cells), MGUS (n=262 cells), and MM (n=2 060 cells). (E) UMAP visualization of representative canonical markers for EC. (F) UMAP projection showing the distribution of MSC from control (n=132 cells), MGUS (n=143 cells), and MM (n=229 cells). (G) UMAP visualization of representative canonical markers for MSC.

To better understand the transcriptional shifts of MSC and EC during MM progression, these two cell populations were analyzed separately (**Fig. 1D-G, Supplementary Fig. 2G-I)**. Overall, these results support our cell enrichment strategy for isolating EC and MSC within the non-hematopoietic BME.

### Identification of a MM-specific IFN signature in the BME

Delving into the composition of EC, our analysis revealed a cluster of cells shared across all stages alongside a second cluster (“MM2”) that was nearly unique to the MM stage (**Fig. 2A**). Cells within the MM2 cluster originated from both male and female mice, indicating that its emergence is not driven by a single animal (**Supplementary Fig. 3**). The “MM2” cluster was characterized by the expression of IFN-inducible genes such as *Ifit2*, *Rsad2*, *Ifit3*, *Isg15*, *Ifi44, Ifit,* and *Oasl2*, indicating an upregulation of an IFN response, as supported by GO ORA (**Fig. 2B,C**).

**Fig. 2.**
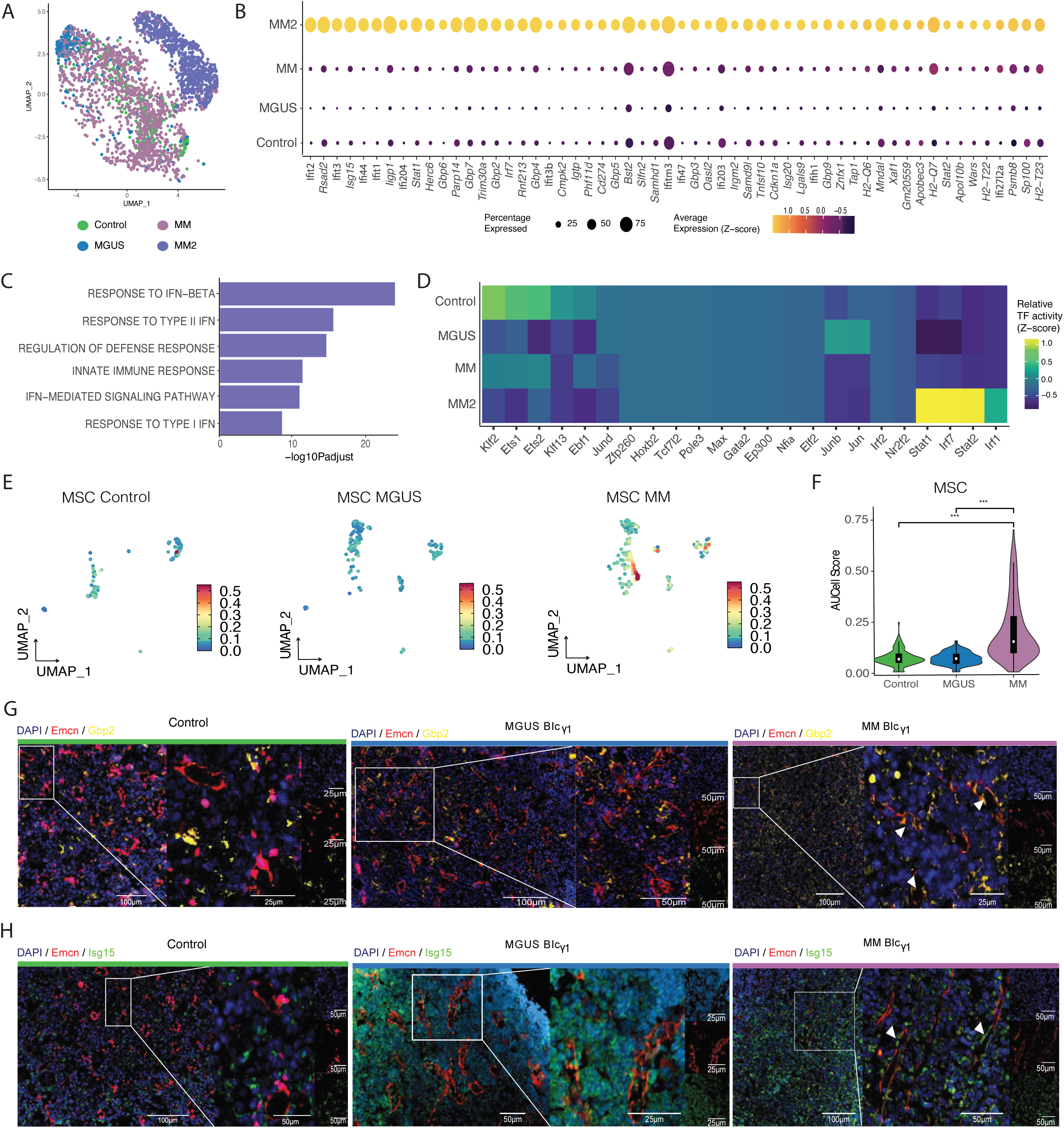
Identification of a MM-specific IFN signature in the BME. (A) UMAP dimension reduction of EC revealing a transcriptionally distinct population in MM. (B) Dot plot visualization of the markers defining EC MM2 cluster. Average gene expression is shown by color intensity, and the percentage of cells expressing each gene is shown by size. (C) Bar charts of enriched GO terms from ORA (adjusted p. value) computed using MM2 cluster markers. The horizontal axis represents the negative of the base 10 logarithm of the adjusted p. value. (D) Heatmap representing z-scaled mean regulon activity score of inferred top specific regulons for each stage. (E) UMAP plot illustrating the distribution of the IFN-related signature in MSC across control, MGUS, and MM. (F) Violin plots showing AUC-score for IFN-related signature within MSC in control, MGUS and MM cells. (G-H) Immunofluorescence (IF) staining of EC (Emcn, red), IFN response genes markers of the MM2 cluster (Gbp2, yellow (G); Isg15, green (H)), and nucleus (DAPI) (blue) in Formalin-Fixed Paraffin-Embedded (FFPE) femurs from Control (YFP_cγ1_), MGUS BI_cγ1_ and MM BI_cγ1_. White arrows point EC Ecmn^+^ IFN^+^ (Gbp2 (G)/Isg15 (H)) in MM BI_cγ1_.

To uncover the putative transcription factors (TFs) that might be underlying the IFN-like signature, we performed GRN inference using SCENIC^16^. The study identified Irf7, Stat1, and Stat2 as potential master regulators significantly activated in “MM2” EC (**Fig. 2D, Supplementary** Fig. 4A-C). These TFs are known positive regulators of type I and type II IFN responses, which are associated with T cell immunosuppressive function in MM patients^17^.

To determine whether the IFN-like signature was unique for EC, we evaluated the signature score related to “MM2” EC across other BM populations. This signature was also significantly upregulated in MM MSC at the MM stage as well as PC, neutrophils, and erythroblasts, suggesting a shared immunomodulatory response of the BME to the disease (**Fig. 2E,F, Supplementary Fig. 4D**). As the signature was not present in control YFP_cγ1_ mice or at precursor stages, we termed it MM-IFN signature to emphasize its association with the myeloma stage. IF staining of BM samples from control YFP_cγ1_ and BI_cγ1_ mice at MGUS and MM stages, validated the increased proportion of IFN^+^ EC and MSC in the MM stage (**Fig. 2G,H, Supplementary Fig. 4E)**.

Interestingly, when the MM-IFN signature was analyzed in the MI_cγ1_ model^13^, which carries MYC and IKK2^NF-κB^ from germinal-center B cells, no significant enrichment at any stage in either population was observed (**Supplementary Fig. 4F-I).** This finding prompted us to investigate this MM-IFN signature in patients. The analysis of the only publicly available dataset from MM patients^11^ that contains non-immune BME uncovered an enrichment of the murine MM-IFN signature in both EC and MSC from a subset of MM patients (**Supplementary Fig. 4I-M**). Likewise, previous studies have described a similar IFN-inducible signature with highly overlapping marker genes in PC and immune microenvironment cells from certain MM patients^3,4^. Collectively, our findings suggest that IFN-signaling is a coordinated response within the BME that could define a subgroup of MM patients.

### A stress pre-vascular state characterizes EC during the MGUS stage

To investigate deeper the molecular mechanisms of myeloma progression, we analyzed the transcriptional changes among EC at the different stages of the disease. An initial analysis uncovered a distinct transcriptome profile of EC at the MGUS stage (**Fig. 3A**). MGUS EC overexpressed *Slpi*, *Hspa5*, *Mt1* and *Mt2*, indicating a stress state at the onset of the disease related to protein folding, reticulum stress, and ion homeostasis (**Fig. 3B, Supplementary Fig. 5A,B, Supplementary Table 3**). The upregulation of these genes may represent a “pre-vascular phase” in the progression to MM. Moreover, *Cd74*, *Tmem176a,* and *Nme2*, were also overexpressed in MGUS EC (**Supplementary Fig. 5C**). DEA between MGUS and non-IFN MM EC (n=1 193 cells) identified 180 differentially expressed genes (DEGs) most of them upregulated (175 upregulated and 5 downregulated) (**Fig. 3C; Supplementary Table 4**), with MM EC showing a pro-vascular and lipid metabolism profile (**Fig. 3D**). *Sox18*, a well described gene involved in tumor vascular development^18^, as well as *Fabp4* and *Cav1* were upregulated, suggesting their role in malignant transformation (**Supplementary Fig. 5D**). In the MGUS-like stage of the MIcγ1 model, we observed a stress-related transitional similar to that seen in BIcγ1. However, MGUS MIcγ1 EC overexpressed genes of MM EC, indicating a stronger vascular phenotype and faster progression (**Supplementary Fig. 6).**

**Fig. 3.**
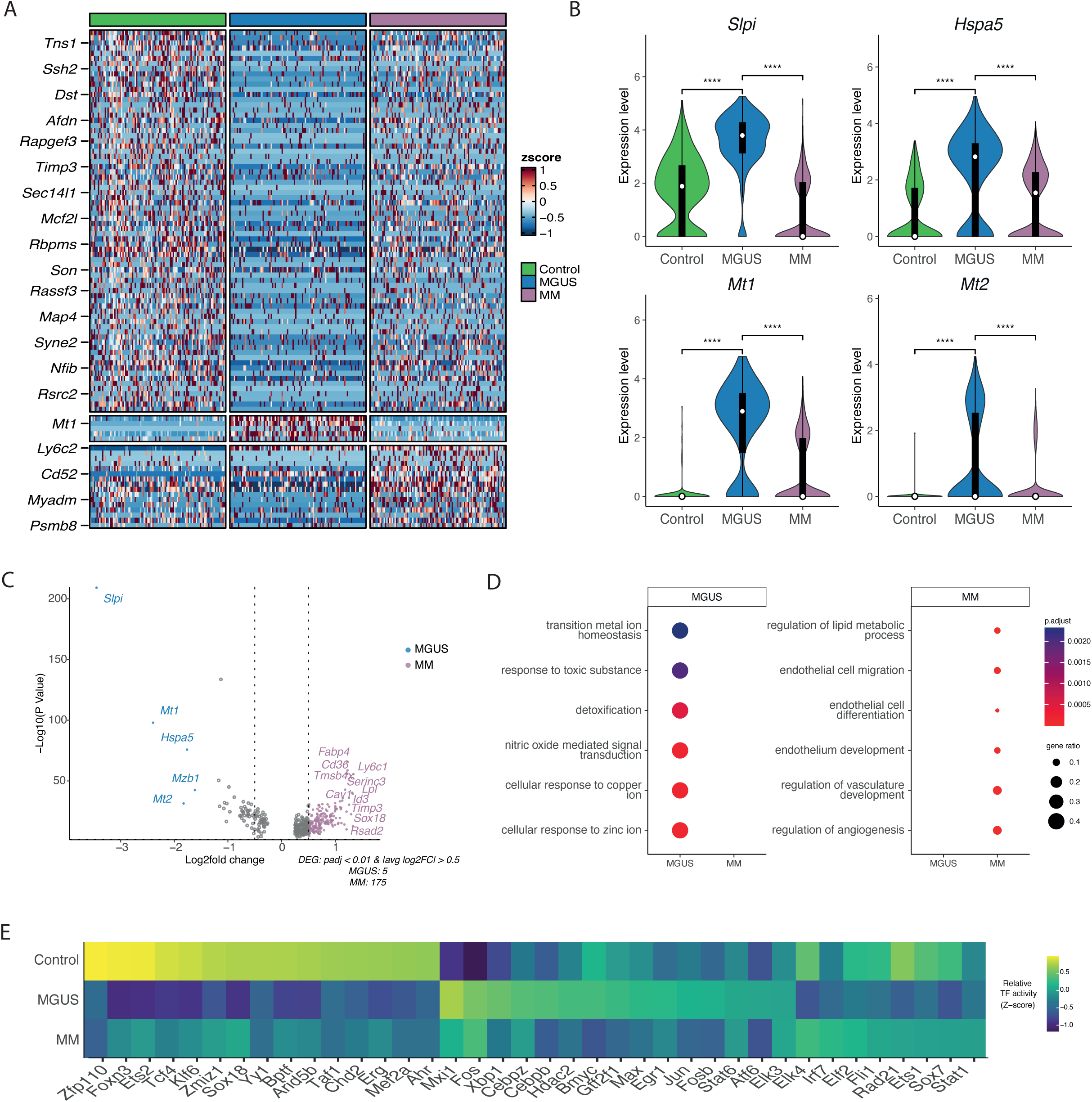
A stress pre-vascular state characterizes EC during the MGUS stage. (A) Heatmap of the DEGs between control, MGUS, and MM cells. Color represents the z-score scaled expression values. (B) Violin plots showing the distribution of expression of genes upregulated in MGUS EC. (C) Volcano plot of the DEGs in MGUS vs. MM EC. The y-axis represents the -log10(p-value), and the x-axis represents the log2(fold change). The dot’s color denotes the disease stage for which DEG was detected, with grey dots representing non-significant genes. (D) Significant terms derived from ORA between EC from MGUS and MM. (E) Heatmap representing z-scaled mean regulon activity score of inferred top specific regulons for EC in each stage.

Next, we identified and examined regulons of the BI_cγ1_ model between MGUS and non-IFN MM EC. The analysis revealed a decline in the activity of regulons during disease progression (**Fig. 3E, Supplementary Fig. 7A,B**). Xbp1, which promotes endothelial proliferation via VEGF ^19^, showed the highest regulon activity in MGUS EC, while Fos regulon was active at both stages. While Fos role has been previously described in PC ^20^ our findings point to a potential regulatory role in EC as well (**Supplementary Fig. 7C,D**). Globally, these findings suggest that MM-associated vascularization is preceded by a stress-related regulatory state in MGUS EC, potentially contributing to disease progression.

### MSC undergo early transcriptional changes during MGUS-like stage that persist in symptomatic MM

Unlike EC, DEA indicated that MSC deregulation begins as early as the premalignant MGUS stage (**Fig. 4A**). GSEA revealed impaired osteogenic and adipogenic differentiation capacity in MSC during MM development, consistent with prior studies^8,10,21^(**Fig. 4B, Supplementary Table 5**). Additionally, we identified downregulation of muscle development genes, including *Pi16*, *Myl6*, *Enpp2, Nupr1*, and *Shox2*, which were not previously linked to MM.

**Fig. 4.**
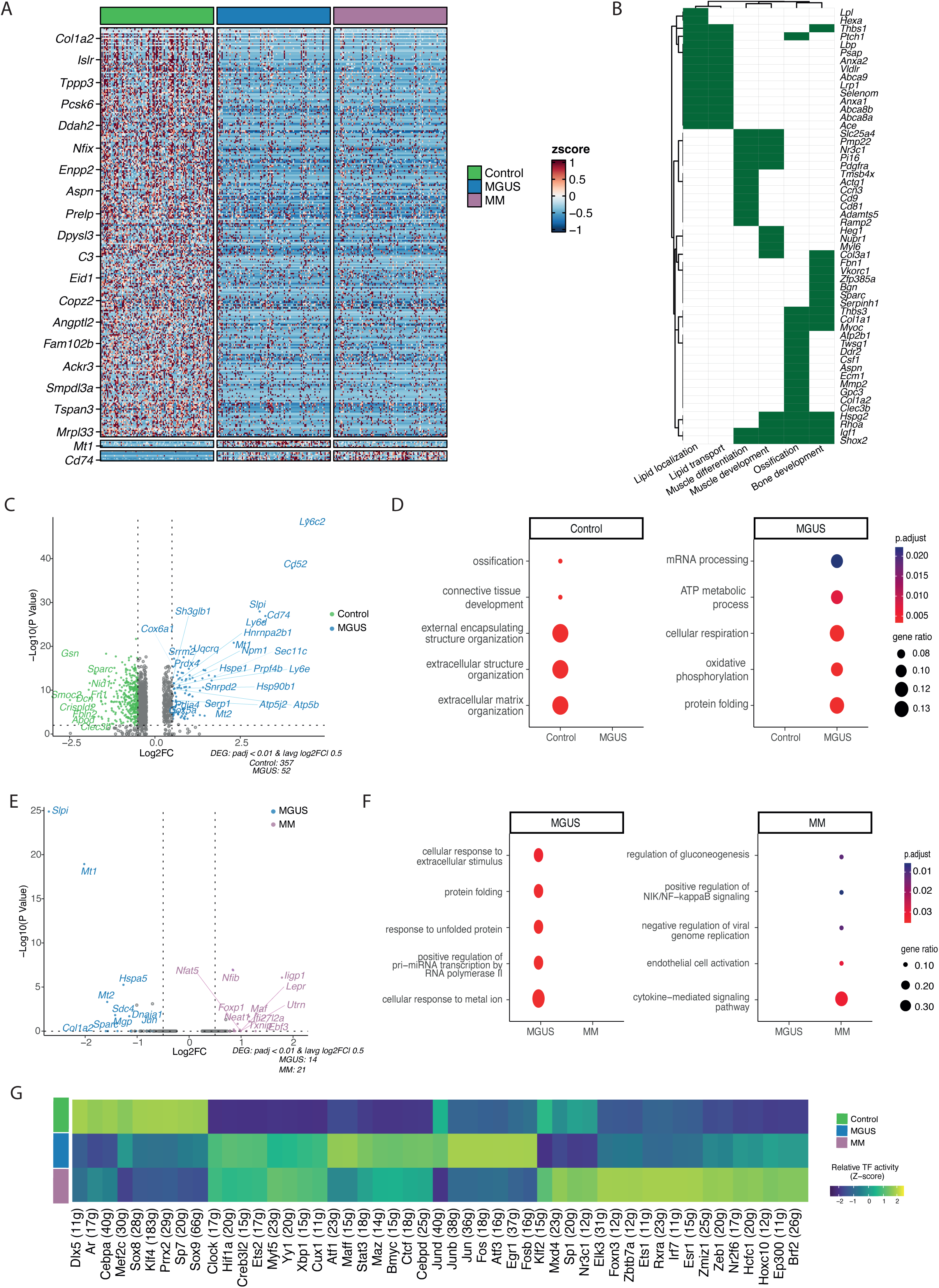
MSC undergo early transcriptional changes during pre-malignant MGUS that persist in symptomatic MM. (A) Heatmap depicting DEGs between MSC in control, MGUS, and MM. Color represents the z-score scaled expression values. (B) Heatmap of significantly downregulated GO terms in MGUS and MM cells compared to control MSC with the associated genes. The green color represents the presence of the gene in each GO term. (C) Volcano plot of the DEGs from the comparison between MSC from control and MGUS. The y-axis represents the -log10(p-value), and the x-axis represents the gene’s log2(fold change). The dot’s color denotes the disease stage for which DEG was detected, with grey dots representing non-significant genes. (D) Significant terms derived from ORA of MSC from control vs MGUS. (E) Volcano plot of the DEGs in MSC from MGUS vs. MM. (F) Significant terms derived from ORA between MSC from MGUS and MM. (G) Heatmap of mean regulon activity score of significantly enriched regulons in each stage. The color scale represents the z-score of the activity scaled by row.

Given early changes at the MGUS stage, we compared MGUS and control MSC, identifying 409 DEGs (52 upregulated and 357 downregulated) (**Supplementary Table 6**). MGUS MSC exhibited enhanced expression of genes and pathways related to oxidative phosphorylation, ATP metabolism, protein folding, and RNA processing (**Fig. 4C,D**). Upregulation of *Cd74* in MGUS MSC and EC suggests its potential role in MM progression. Only 35 DEGs were identified between MM and MGUS MSC (21 upregulated and 14 downregulated), further reflecting the high level of transcriptional similarity between both stages. Upregulated genes in MM MSC were mainly related to immune response, consistent with the inflammatory profile in human MM MSC^11^ (**Fig. 4E,F, Supplementary Table 7).** Similar transcriptional changes were observed in MIcγ1 mice, supporting the impairment of MSC differentiation early in the MGUS-like stage (**Supplementary Fig. 8**).

Finally, a regulon analysis identified key TFs driving MSC alterations during MM progression (**Fig. 4G, Supplementary Fig. 9A**). In control MSC Sp7, Dlx5, Sox8, Sox9, and Cebpa, well-known TFs mediating osteochondrogenic and adipogenic differentiation, were identified. Reduced Klf4 activity supported impaired myogenic potential in MGUS and MM MSC. Moreover, TFs from the AP-1 family and Egr1 showed the highest regulon activity in MGUS MSC, consistent with their implication in MM development^22,23^. Yy1 and Xbp1 regulons remained active from MGUS to MM, while Elk3 was specific to MM, suggesting its role in cancer cell invasion^24–26^ (**Supplementary Fig. 9B,C**). Overall, our results suggest that MSC undergo a TF-mediated transcriptional shift at the MGUS stage that persists through disease progression.

### Altered crosstalk between EC and MSC with malignant PC is potentially associated with myeloma development

To further understand the mechanisms underlying BME-PC communication and how they may change upon disease progression, we analyzed cellular communication through LIANA ligand-receptor (L-R) analysis^27^ (**Fig. 5A, Supplementary Fig. 10, Supplementary Table 8**). Interactions, through the *Fn1* ligand and *Insr* receptor in EC and the *Cd93* PC receptor, were identified only in controls, while *Apoe* ligand in EC and *Adam9*, *C3,* and *Vegfa* PC ligands were only found in MM (**Fig. 5B**). Our results support the role of angiogenesis in myeloma via *Vegfa* secretion from PC, interacting with *Kdr*, *Nrp1*, and *Nrp2* in EC. High expression of Nrps has been associated with tumor angiogenesis^28^. In addition, *Efna1*-*Epha8* interaction was predicted exclusively in MM.

**Fig. 5.**
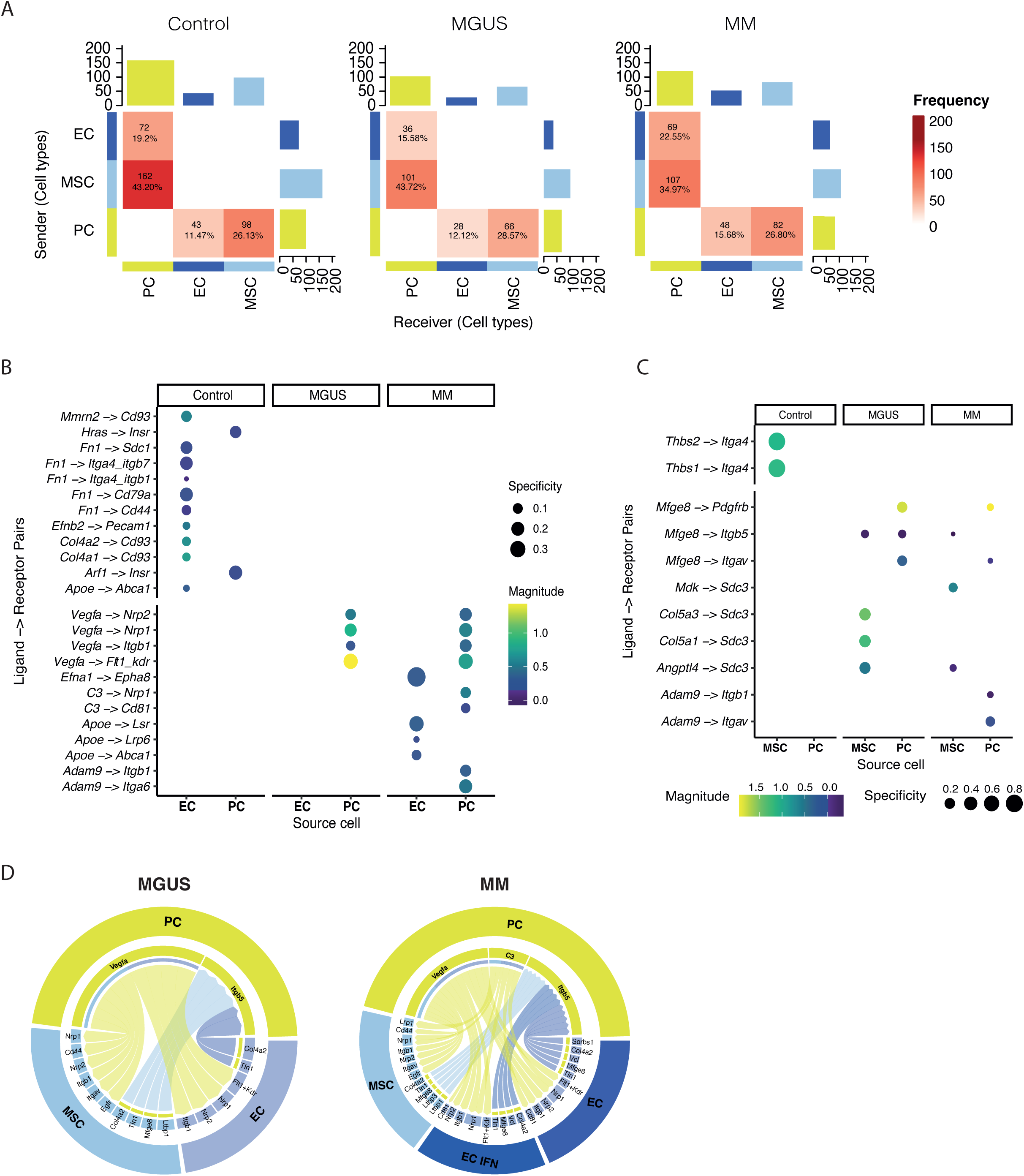
Altered crosstalk between EC and MSC with malignant PC is potentially associated with myeloma development. (A) Heatmap showing the number of predicted cellular interactions between EC, MSC and PC. Block sizes and colors are proportional to the number of ligand-receptor pairs. (B-C) Dot plots of representative ligand-receptor interactions between PC and EC (B) or MSC (C) during disease progression. Dot size and color indicate the interaction’s specificity and strength, respectively. (D) Chord diagram showing a shared relevant signaling network between EC, MSC, and malignant PC in the MGUS and MM stages. Colors and width represent signal senders and strength, respectively.

Interestingly, analysis of MSC-PC communication revealed interactions related to cell motility and adhesion in MM that may be related to malignant PC extravasation (**Supplementary Fig. 10D)**. We observed a decrease of *Thbs1-Itga4* and *Thbs2-Itga4* interactions, whereas MGUS and MM PC participate in outgoing interactions involving *Mfge8* and *Sdc3*-mediated incoming interactions (**Fig. 5C**). Likewise, *Adam9,* which promotes cancer egress and invasion^29^, interacts with MSC through *Itgav* and *Itgb1* in MM.

We uncovered common L-R pairs between EC-PC and MSC-PC along disease progression (**Fig. 5D**). Multiple interactions through the *Itgb5* PC receptor were initiated during MGUS. Moreover, in line with the previous study in human MM patients^11^, malignant PC interacted with MSC and EC through *Vegfa* secretion, highlighting the role of angiogenesis in MM. Additionally, *C3*-mediated interactions between PC and the non-hematopoietic BME in MM suggest that *C3* crosstalk maintains the immunosuppressive environment. Overall, despite the limitations associated with the sparsity of single-cell resolution data and the limited number of cells analyzed, the BME-PC interactions identified in the L-R based analysis are consistent with known pathways associated with MM further identifying potential targets in the disease.

### Treatment of MM mice with VRd induces a transcriptional remodeling of EC and MSC

To assess the impact of MM therapy on EC and MSC molecular signatures, we performed scRNA-seq on a pool of BI_cγ1_ mice carrying humanized CRBN^I391V^ (BI_cγ1_-*Crbn*^I391V^)^14^ after 20 weeks of treatment with VRd, a current standard of care for patients with MM (**Supplementary Fig. 11A-D**). As expected, VRd treatment significantly reduced the tumor burden and improved survival outcomes (**Supplementary Fig. 11E,F**). Given the role of IFN signaling as a myeloma-specific response, we first sought to determine the presence of IFN signature after treatment. Interestingly, VRd therapy downregulated IFN-driven MM signaling in EC and other BM populations (**Fig. 6A, Supplementary Fig. 11G**). As a result, EC from treated mice exhibited decreased expression of IFN-induced genes, such as *Ifit1, Ifit2, Gbp2*, and *Isg15*, and key related regulons previously upregulated during myeloma development (**Supplementary** Fig. 11H**, Supplementary Table 9**). IF analysis further confirmed the decrease in IFN^+^ EC and MSC following VRd treatment, supporting a potential link between IFN signaling suppression and the anti-myeloma drug response (**Fig. 6B,C, Supplementary Fig. 11I)**. To further explore the significance of our findings, we analyzed a publicly available bulk RNA-seq data of MSC from MM patient samples before and after induction therapy with bortezomib-thalidomide-dexamethasone with or without daratumumab^11^. This revealed a reduction of IFN-signature post-treatment in high-IFN MM patients (**Supplementary Fig. 11J,K)**. Similarly, Chen et al.^6^ reported a widespread downregulation of IFN-response genes in immune cells following bortezomib-cyclophosphamide-dexamethasone therapy, supporting the relevance of our results in the context of human disease.

**Fig. 6.**
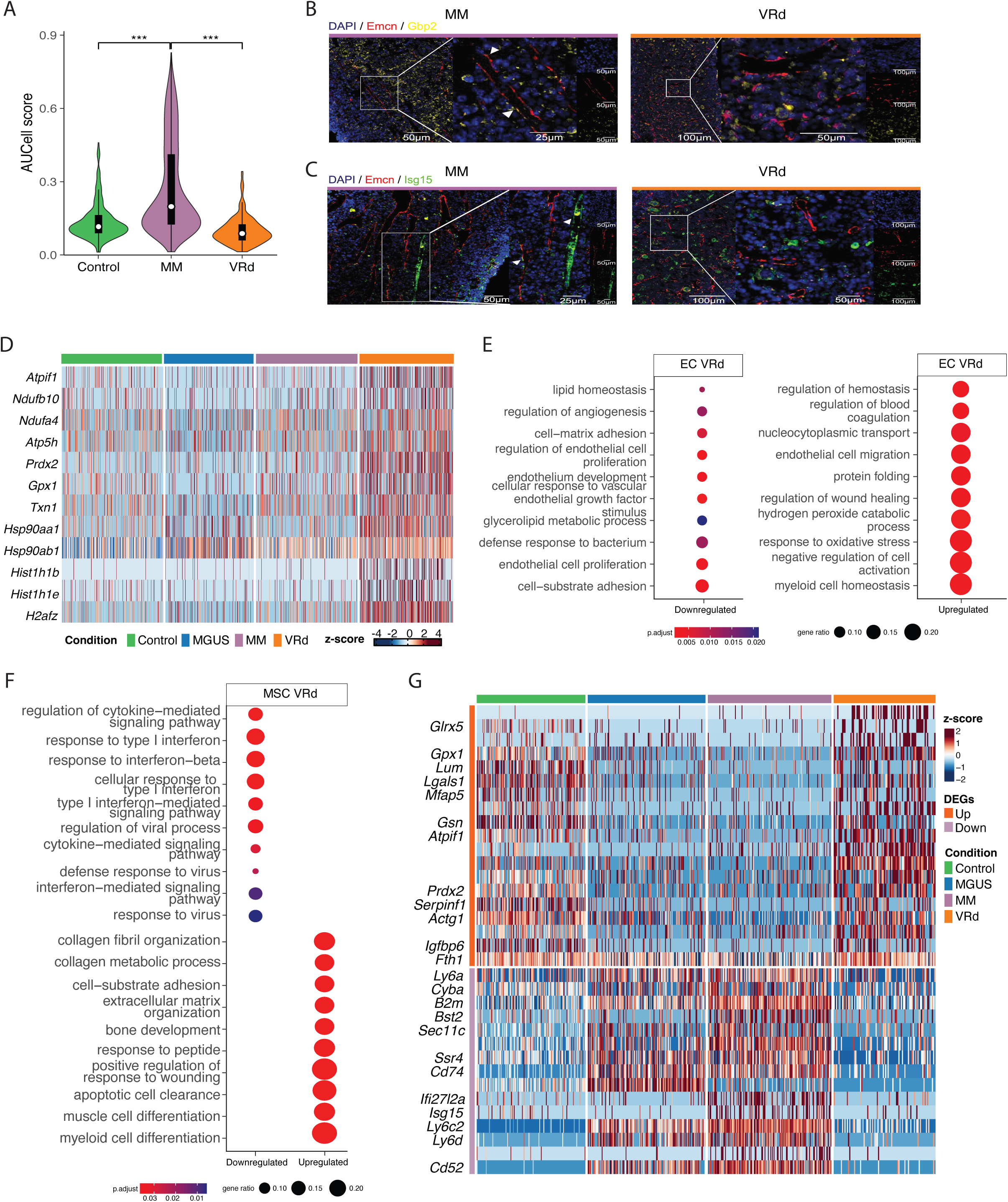
scRNA-seq reveals VRd-induced EC and MSC remodeling in myeloma mice. (A) Violin plot showing AUC-score for IFN-related signature in EC from Control (YFP_cγ1_), MM BIc_γ1_ (MM) and MM BI_cγ1_-*Crbn*^I391V^ VRd-treated (VRd) mice. (B-C) IF staining of EC (Emcn, red), IFN response genes (Gbp2 (yellow, (B)); Isg15 (green, (C)), and nucleus (DAPI) (blue) in FFPE femurs from MM BI_cγ1_ mice (MM) and VRd-treated mice. White arrows point EC Ecmn^+^ IFN^+^ (Gbp2 (B)/Isg15 (C)) in MM BI_cγ1_. (D) Heatmap showing the expression of stress-related DEGs identified in the comparison between treated and non-treated non-IFN MM EC at different stages of the disease. Color represents the z-score scaled expression values. (E) Significant terms derived from ORA between VRd-treated and untreated non-IFN MM EC. (F) Significant terms derived from ORA of MSC from MM mice before and after VRd treatment. (G) Heatmap projecting the expression of DEGs identified in the comparison between non-treated and treated MSC at different stages of the disease. Color represents the z-score scaled expression values.

Beyond dampening the IFN signature, VRd treatment induced broader gene expression changes in both EC and MSC. In EC, DEA revealed upregulation of genes involved in mitochondrial metabolism, stress response, detoxification, and chromatin remodeling—indicative of an adaptive response (**Fig. 6D,E)**. Consistent with its known tumor-suppressive role and previous studies in MM cells after bortezomib exposure, *Klf2* was upregulated in VRd-treated EC^30,31^ (**Supplementary Fig. 11L)**. Furthermore, GRN analysis revealed enhanced activity of Zbtb7a and Rarg regulons, reinforcing this adaptive response^32,33^ (**Supplementary Fig. 12A)**. Treated EC were enriched in thrombotic regulation pathways, including upregulation of *Hmgb1*, a gene linked to MM drug resistance^34^ (**Supplementary Fig. 12B**). Consistent with lenalidomide’s anti-angiogenic effects^35,36^, vascular and adhesion-related genes, including *Nrp1*, were downregulated and Xbp1 regulon activity was reduced (**Supplementary Fig. 12A)**.

In MSC, VRd treatment downregulated immune and IFN signaling pathways, indicating reduced inflammation in the BME (**Fig. 6F)**. Likewise, antigen presentation genes, such as *Cd74*, *B2m* and *Psmb8* were downregulated (**Fig. 6G, Supplementary Table 10)**. Interestingly, decreased expression of *Mdm4* and *Cyba* in treated MSC suggests that VRd may impair the proliferative supportive capacity of MSC^37,38^ (**Supplementary Fig. 12C)**. On the contrary, genes linked to ECM organization, wound healing, and bone development were upregulated, aligning with bortezomib’s anabolic effects^39,40^ **(Supplementary Fig. 12D).** GRN analysis confirmed osteogenic reprogramming via reactivation of Cebpa, Nfib, Nfic, Churc1 and Sox4 regulons (**Supplementary Fig. 12E)**. Finally, MSC-PC communication analysis identified interactions previously observed in control MSC such as *Adam10–Tspan5*, *Adam17–Il6ra* and *Wnt10a–Fzd1/Lrp6* which may contribute to anti-inflammatory and osteogenic adaptations following VRd treatment^41–44^ (**Supplementary Fig. 12F)**. Overall, our results suggest that VRd therapy suppresses IFN-driven MM signaling and reprograms the BM niche by triggering an adaptive response in EC and restoring MSC osteogenic potential, creating a less tumor-permissive BME.

### Transcriptional dynamics of human MSC during myeloma progression

To validate our findings in mice, we performed bulk RNA-seq in FACS-isolated MSC from aged-matched HD (n = 8), newly diagnosed MGUS (n = 5), SMM (n =2), or MM (n =10) patients (**Fig. 7A, Supplementary Fig. 13A**). PCA revealed a distinction between MM-MSC and HD-MSC, with MGUS-MSC and SMM-MSC mapped between them, showing an evolving transcriptional trajectory (**Fig. 7B**).

**Fig. 7.**
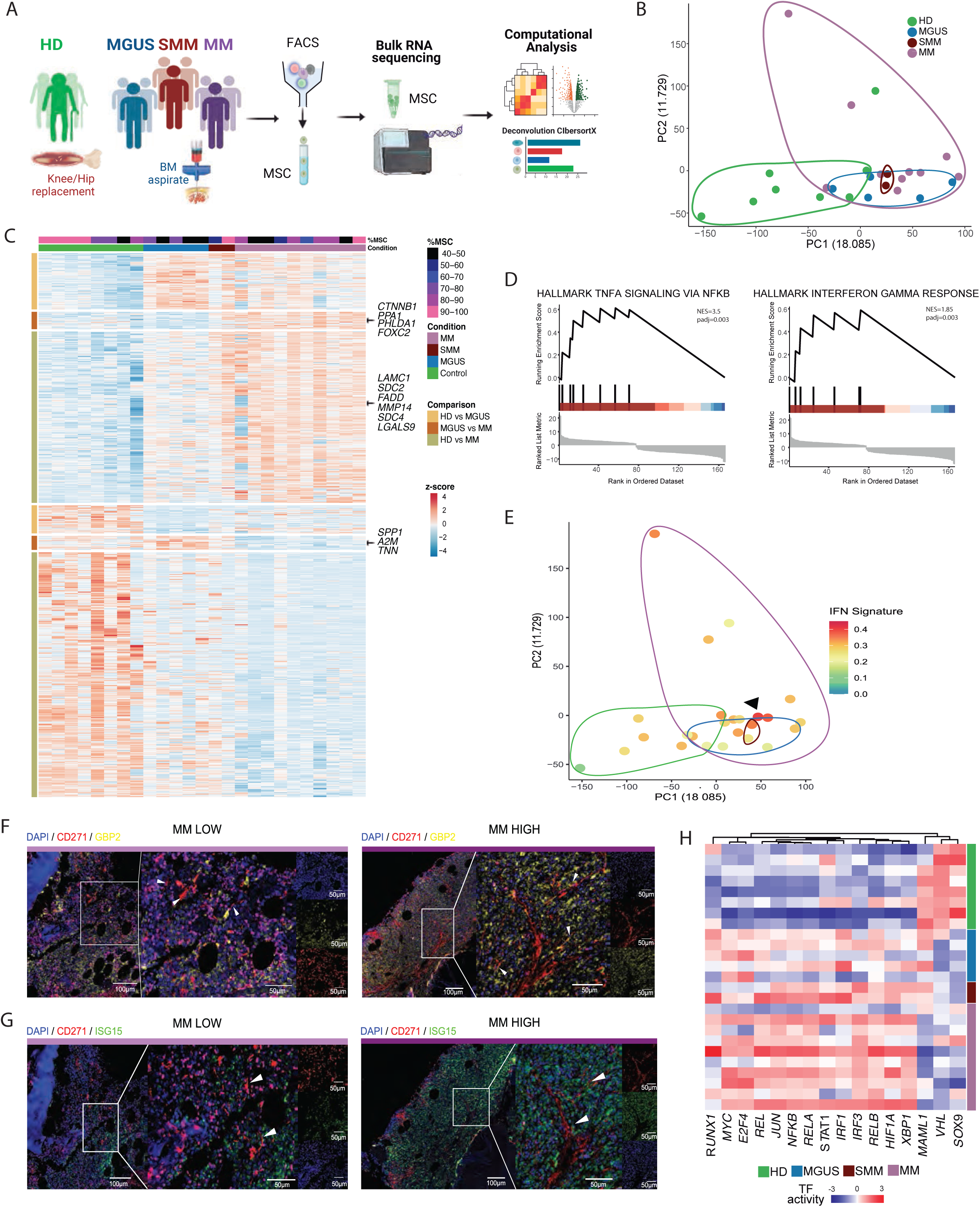
Transcriptional dynamics of human MSC during myeloma progression. (A) Experimental workflow used for the isolation and bulk RNA-seq analysis of human BM MSC. (B) PCA of MSC expression profiles shows clustering by disease stage. The color of the dots represents disease stages. (C) Heatmap showing the expression profiles across samples of DEGs identified in each pairwise comparison along disease progression. (D) GSEA plots for the gene sets ‘TNFα signaling via NFκB’ (left panel) and ‘Interferon gamma response’ (right panel) in the comparison between healthy donors (HD)-MSC and MM-MSC. (E) PCA plot showing the MM-IFN signature score in each human sample, previously identified murine BM MM cells. (F-G) IF staining of MSC (CD271, red), IFN response genes (*GBP2*, yellow (F); *ISG15,* green (G)), and nucleus (DAPI) (blue) in archived paraffined human BM biopsies of MM patients with low (MM LOW) and high IFN signal (MM HIGH) in stromal cells (see Supplementary Table 2). White arrows point MSC CD271^+^ in MM LOW IFN samples, and MSC CD271^+^ IFN^+^ (GBP2/ISG15) in MM HIGH IFN samples. (H) Heatmap representing the variation of TF activity score across samples.

The comparison between MGUS-MSC and HD-MSC revealed 180 DEGs (120 upregulated and 60 downregulated) (**Fig. 7C, Supplementary Fig. 13B, Supplementary Table 11**). Downregulation of genes related to osteoblast differentiation, such as *TNN, SPP1,* and *A2M,* indicate the early remodeling of MSC previously described in our mouse data. Besides, MGUS-MSC upregulated genes related to cell cycle, protein ubiquitination, ATP metabolism, and oxidative stress supporting their role in MM pathogenesis (**Supplementary Fig. 13C**).

Consistent with the high transcriptional similarity observed in mice between MGUS and MM MSC, the fewest DEGs were yielded when comparing MGUS and MM-MSC (41 upregulated and 33 downregulated) (**Fig. 7C, Supplementary Fig. 13D, Supplementary Table 12**). MM-MSC showed enrichment in inflammatory-related processes, indicating that a pro-inflammatory microenvironment is required for the transition between premalignant stages and MM, in line with previous studies^12,45^. In addition, MM-MSC over-expressed *PHLDA1*, *FOXC2*, *PPA1*, and *CTNNB1,* genes associated with Epithelial-Mesenchymal Transition (EMT), which aligns with MSC potential to induce EMT through extracellular vesicles in MM^46^ (**Supplementary Fig. 13D,E)**.

Defective osteogenic differentiation and dysfunctional immunomodulatory ability were confirmed when comparing MM-vs HD-MSC (448 upregulated and 566 downregulated) (**Fig. 7C, Supplementary Fig. 13F,G, Supplementary Table 13)**. Furthermore, consistent with the impaired myogenic phenotype observed in mice, human MM-MSC showed downregulation of muscle-related genes. GSEA revealed enrichment of TNF-α signaling via NF-ΚB and IFN response in MM-MSC (**Fig. 7D)**. Interestingly, the IFN signature previously identified in murine MM BME was enriched in a subset of MM patients, as shown in the analysis of publicly available data from de Jong et al.^45^ (**Fig. 7E, Supplementary Fig. 4I-L, Supplementary Fig. 11J,K, Supplementary Fig. 13H**). IF staining of BM samples from MGUS and MM patients showed an enrichment of IFN-driven MM signaling in MSC from a subset of MM patients, further supporting our transcriptional findings (**Fig. 7F,G, Supplementary Fig. 14A,B)**.

Finally, we determined changes in TF activity during myeloma progression (**Fig. 7H**). Enhanced activity of TFs related to the NF-ΚB pathway, such as NF-ΚB, NKFB1, and RELA, were observed in MSC from both MGUS and MM patients, highlighting NF-κB activation as an early event in MM evolution. Furthermore, MM-MSC exhibited increased activity of IRF1 and IRF3, which again emphasizes the importance of IFN signaling for the disease. In conclusion, the transcriptional changes of MSC between MSC from healthy donors, MGUS and MM patients mirrors our findings in mouse models.

## DISCUSSION

Over the last decades, our understanding of MM biology has advanced significantly ^47–51^, explicitly revealing the contribution of the BME to the outgrowth and relapse of this hematological cancer and exerting a supportive function for the expansion of malignant PC^3,11,52–54^. However, research has mainly concentrated on PC and their immune microenvironment, highlighting the need for additional exploration of the non-hematopoietic BME^9,55–57^. Here, using our validated BI_cγ1_ and MI_cγ1_ murine models^13^, that closely mimic key features of human MM progression, we provided a comprehensive overview of transcriptional deregulation in EC and MSC during the transition from healthy to MGUS and MM as well as the reshaping of the non-hematopoietic BME in response to VRd treatment. Further, we validated our results in MSC from MM patients at various disease stages, both in our own data as well as in the only publicly available single cell MM patient dataset^11^.

Through analysis of the transcriptomic alterations along disease progression, we uncovered a pre-vascular state in MGUS EC linked to protein folding and ion homeostasis, where upregulation of *Slpi*, *Mt1*, *Mt2*, and *Hspa5*, could potentially serve as an early myeloma fingerprint in EC. Disease progression from MGUS to MM in EC involved angiogenesis, migration, and metabolic shifts. Xbp1 and Fos, key regulators of vascular development and cancer progression^19,20^, showed enhanced activity with myeloma progression highlighting their potential as disease targets. In contrast, transcriptional changes in MSC previously described in advanced MM patients^8,10^, were already present during the precursor MGUS-like stage. In addition to impaired osteogenic differentiation, MGUS MSC exhibited a downregulation of myogenesis-related genes. Consistent with these murine findings, our bulk RNA-seq showed impaired MSC differentiation in human patients, even at premalignant stages. Similarly, MGUS prompted a metabolic swift in human MSC with increased protein folding. These findings suggest that MGUS MSC show a reshaping of their differentiation capacity and metabolism to support the growth of MM cells from the early stages of the disease.

Understanding the crosstalk between the BME and PC is crucial for unraveling tumor progression mechanisms, and this study contributes to understand the interactions between EC, MSC, and malignant PC along the disease evolution. Consistent with a previous study in MM patients^11^, we found *Vegfa* secretion from PC as a hallmark of MM angiogenesis through the interaction with *Kdr*, *Nrp1*, and *Nrp2* endothelial receptors. Additionally, identifying *Efna1-Epha8* interaction exclusively in MM implicates the ephrin pathway in disease development, as previously described in other cancers^58^. MSC-PC interactions were linked to adhesion, matrix organization, and migration, suggesting MSC-PC crosstalk is crucial for MM extravasation and disease dissemination.

A fundamental finding of our study is the identification of a myeloma-specific IFN signature in EC that extends to MSC and other BM cells, indicating a shared BME response during MM progression. This signature, that differs from the inflammatory phenotype via TNFα and NF-κB described by de Jong et al.^11^, is marked by upregulation of IFN-inducible genes and master regulators (Irf7, Stat1, Stat2). STAT1/2 signaling, closely linked to IFN in MM, supports tumor-promoting roles, complementing *STAT3*’s known oncogenic function in MM migration and BM homing^59^. While prior scRNA-seq studies reported IFN activity in PC and immune cells^3,4^, only limited evidence suggested an IFNα response in MSC^9,55^. The MM IFN-inducible response uncovered in our study, reveals an immunomodulatory role of the non-hematopoietic BME, with EC and MSC potentially contributing to immune remodeling and T-cell exhaustion under inflammatory conditions^17^.

Our findings expand on the concept of pro-inflammatory signaling within MM MSC^11^, identifying an enrichment of this IFN signature in both EC and MSC from a subset of MM patients, highlighting its clinical relevance for patient stratification based on inflammatory features. More importantly, the suppression of this MM-IFN signature in human MM patients after induction therapy, further supports its clinical relevance. In addition, using our MGUS-MM mouse models, we demonstrate that VRd treatment impacts the non-hematopoietic BME by suppressing this MM IFN signaling. While combinatorial regimens improve outcomes in patients by targeting multiple pathways our results support their potential to reshape the BME^60^ by inducing and adaptive stress in EC, and restoring MSC osteogenic potential, shifting the niche toward a less tumor-permissive state. These findings highlight the importance of targeting BME interactions and enhancing anti-myeloma immune responses.

In summary, the use of the BI_cγ1_ myeloma murine model, which mimics MGUS-to-MM progression, alongside scRNA-seq, has enabled a comprehensive dissection of the non-hematopoietic BME. The myeloma-specific IFN signature, the transition stress state in MGUS EC, the early transcriptional shifts in MSC, the evolving cellular communication networks and the IFN signaling repression post-VRd therapy collectively support the role of EC and MSC in MM development and treatment response. Validation in human patients reinforces the study’s translational relevance, suggesting that non-hematopoietic BME changes may foster MM development from premalignant stages and shift toward a less tumor-permissive state after treatment.

## ACKNOWLEDGEMENTS

This work was supported by the Instituto de Salud Carlos III and co-financed by ERDF A way of making Europe (PI20/01308, PI22/00983, PI23/00516); CIBERONC (CB16/12/00489), RICORS TERAV (RD21/0017/0009); Departamento de Industria Gobierno de Navarra (AGATA 0011-1411-2020-000010/0011-1411-2020-000011) and Departamento de Salud Gobierno de Navarra; the Cancer Research UK [C355/A26819]; FC AECC and AIRC under the Accelerator Award Program; The International Myeloma Foundation (Brian van Novis) and The Paula and Rodger Riney Foundation to FP. The study was also supported by The Spanish Government, through project PID2019-111192GA-I00 (MICINN) to DGC. IAC was supported by Marie Curie grant (H2020-MSCA-IF-837491) from European Commission. IC was supported by AECC predoctoral fellowship (PRDNA19006CENZ). MC was supported by FPU fellowship (FPU22/03283) from Ministerio de Ciencia, Innovación y Universidades.

We would like to thank the staff of the flow cytometry core, the advanced genomic lab, and the animal facility at CIMA Universidad de Navarra for their invaluable technical and intellectual assistance. We also acknowledge Hospital Reina Sofía de Tudela, Hospital Universitario de Navarra, the Biobank of the University of Navarra, and Goretti Ariz for the provision of BM samples and their collaboration. We are particularly grateful to the patients and healthy donors for their participation in this study.

## AUTHORSHIP CONTRIBUTIONS

I.C., M.C., L.C-D. and I.A.C. processed murine and human BM samples; B.S. and I.A.C. conceptualized the study and analyzed the scRNA-seq experiments data; I.C., M.C., and I.A.C. interpreted the scRNA-seq experiments data; I.C. and I.A.C. wrote the paper; M.C. analyzed the bulk RNA-seq experiments data; M.C. and I.A.C. interpreted the bulk RNA-seq experiments data; M.L. and J.A.M.C. designed and provided the mice models; M.L. and S.S. performed the VRd treatment experiments. A.B. performed the cell-cell computational analysis; A.V., P.S.M., and P.A.R. performed the scRNA-seq and bulk RNA-seq experiments. D.A. and A.L. performed the FACS sorting. I.S.G. and J.R. provided the BM samples from HD; M.C., L.C-D. and P.R-C. conducted the histologic experiments; M.C. analyzed the histologic experiments; M.A-M. and E.M. provided the human MGUS and MM samples. D.G.C., with help from M.H. and J.T., supervised the computational work. B.E. provided the humanized mouse model; B.P. and J.S-M. helped with the clinical data; P.R.O. and L.E.T-A. provided the BM samples from MGUS, SMM, and MM patients; I.A.C. and B.S. designed the research experiments; F.P., D.G.C. and I.A.C. conceived and directed the research project.

All authors actively participated in the discussions underlying this manuscript. I.C., M.C., I.A.C., D.G.C., and F.P. discussed the results and wrote the final manuscript. All authors contributed to, read, and approved the final manuscript.

