## Supplementary material for "Stromal and Endothelial Transcriptional Changes during Progression from MGUS to Myeloma and after Treatment Response": LDSupplementalMaterial_Cenzano_BM Niche MGUS-MM_Treatment.docx

SUPPLEMENTAL Methods

Mice

A detailed description of all mice utilized in the study is described in **supplemental Table 1**. We used BI_cγ_ mice (B6(Cg)-Gt (ROSA)26Sor^tm4(Ikbkb)Rsky^/J mice and B6. Cg-Tg(BCL2)22Wehi/J mice crossed with cγ1-cre mice (B6.129P2(Cg)-Ighg1^tm1(cre)Cgn^/J mice; all from The Jackson Laboratory) that recapitulate the most common changes observed in human multiple myeloma (MM) preceded by a monoclonal gammopathy of undetermined significance (MGUS)-like stage^1^. In addition, we used MI_cγ_ mice (B6(Cg)-*Gt (ROSA)26Sor^tm4(Ikbkb)Rsky^*/J and C57BL/6N-Gt(ROSA)26Sor^tm13(CAG-MYC,-CD2*)Rsky^/J mice crossed with cγ1-cre mice), a fast progression model with a less marked MGUS stage. As controls, mice carrying a YFP reporter (B6.129 × 1-Gt (ROSA)26Sor^tm1(EYFP)Cos^/J mice; The Jackson Laboratory) were crossed with cγ1-cre mice. To induce plasma cell (PC) generation in mice kept in specific pathogen-free (SPF) animal facilities, eight to ten-week-old mice were immunized with sheep red blood cells (SRBCs) every 21 days for 3-4 months.

To assess the role of bone marrow microenvironment (BME) in treatment response, we used triplet therapy VRd of bortezomib (Velcade®), lenalidomide (Revlimid®), and dexamethasone on BI_cγ1_, with regimens optimized based on human protocols^2^. Given mouse cells´ inherent resistance to immunomodulatory drugs (IMiDs), BI_cγ1_ mice were crossed with a humanized cereblon (*Crbn* ^I391V^) gene strain, enabling sensitivity to IMiDs^1,3,4^. Before initiating therapy, tumor burden was assessed by quantifying the serum immunoglobulin gamma fraction (M-spikes) by electrophoresis. Treatment was administered upon the emergence of M-spikes. Mice received 1mg/kg (Monday and Friday) of bortezomib (Velcade®), lenalidomide (Revlimid®), 50mg/kg (Monday, Wednesday and Friday), and dexamethasone, 20mg/kg (Monday) along 20 weeks. Animals were monitored twice weekly to detect signs of discomfort and/or disease, included hunching, ruffled fur, labored breathing, low body temperature, low mobility and/or >20% weight loss from the time of study initiation.

For each single cell RNA sequencing (scRNA-seq) experiment, a pool of four, five or six adult mice per group compromising control, MGUS, and MM stages, as well as velcade, lenalidomide and dexamethasone (VRd)-treated subjects, was used. Mice of both sexes were used in the study. Mice were kept under SPF conditions in the Center for Applied Medical Research (CIMA) animal facilities at the University of Navarra. All animal experimentation was approved by the Ethics Committee for Clinical Research at the University of Navarra (protocol CEEA082-20).

Isolation and fluorescence-activated cell sorting of murine BM microenvironment and PC

Mice were euthanized by CO_2_ asphyxiation. Bones from the humerus, radius, iliac crests, femurs, and tibia were harvested in Phosphate-buffered saline (PBS) 1X containing 2 % Fetal bovine serum (FBS) and two mM EDTA (modified PBS). All steps were performed on ice to preserve cell viability and RNA integrity. Muscles and soft tissue were removed from the bones, and bone marrow (BM) cells were obtained by crushing in modified PBS. Cells were then filtered through a 70 μm cell strainer, and red blood cells were lysed with ACK buffer (NH_4_Cl 150 mM, KHCO_3_ 10 mM, and Na_2_EDTA 0,1 mM) for 10 minutes at room temperature (RT). 25 μl were stained with PE-CD138, APC-B220, and BV421-IgM antibodies (Biolegend) and analyzed by flow cytometry to determine PC infiltration (**supplemental Table 1)**. The remaining bone fragments were collected on a 50 ml conical tube, digested with the appropriate volume of PBS (3ml/mouse) with 0.3% collagenase I, and dispase (5U/ml) for 15 min at 37ºC and shaking at 200 rpm. FBS, representing 10% of the digestion volume, was added to stop the digestion of collagenase. After digestion, the collagenized fractions were filtered through a 70 μm filter into a collection tube and pooled into one sample. To determine live cell concentration and viability, 10 μl of each mouse sample was stained with acridine orange and propidium iodide (AO/PI) solution (Nexcelom) and analyzed with Cellometer K2 Image Cytometer (Nexcelom Bioscience). Cells were subsequently stained for 20 minutes on the ice first in the appropriate volume of modified PBS 1X (3 ml/mouse) with 160 μl/mouse of biotinylated lineage cocktail (Mac1, CD3, Gr1, B220, and TER119) followed by incubation with streptavidin magnetic microbeads (100 μl/mouse). Negative selection was performed using Miltenyi LD columns according to the manufacturer's protocol, and resulting cells from both fractions were counted as described before to confirm selection efficiency.

After selection, the negative fraction was stained with the following combination of conjugated antibodies at a concentration of 1/200: APC-Cy7 labeled streptavidin, BV510 labeled anti-CD45, APC labeled anti-CD45, and APC labeled anti-Ter119. Samples were then stained with 0.05 μM of Vybrant dye orange (VDO) at 37ºC for 30 minutes to label living cells. 7AAD was added to discard apoptotic and dead cells from the sample. Cells were stained with one μl/mouse of Annexin V-FITC for annexin V staining on an appropriate volume of 1X Annexin V binding buffer in the dark for 15 min at RT. Samples were resuspended in 1X Annexin V buffer, and five μl of 7AAD dye (up to 1x10^6^ cells) were added.

BM non-hematopoietic cells and GFP-positive (GFP^+^) PC were FACS using a BD FACSAria II sorter and collected in PBS 1X supplemented with 0.05% UltraPure BSA for the subsequent sequencing protocol. Cell viability of sorted cells was assessed using a Nexcelom Cellometer, as described above.

Single cell RNA sequencing

scRNA-seq was performed using the Single Cell 3’ Reagent Kits v3.1 (10X Genomics) according to the manufacturer’s instructions. PC and BME sorted cells for each sample were pooled before scRNA-seq was performed. Approximately 15,000 cells were loaded at a concentration of 1,000 cells/µL on a Chromium Controller instrument (10X Genomics) to generate single-cell gel bead-in-emulsions (GEMs). In this step, each cell was encapsulated with primers containing a fixed Illumina Read one sequence, followed by a cell-identifying 16 bp 10X barcode, a 12 bp Unique Molecular Identifier (UMI), and a poly-dT sequence. A subsequent reverse transcription yielded full-length, barcoded cDNA. This cDNA was then released from the GEMs, PCR-amplified, and purified with magnetic beads (SPRIselect, Beckman Coulter). Enzymatic Fragmentation and Size Selection was used to optimize cDNA size before library construction. Illumina adaptor sequences were added, and the resulting library was amplified via end repair, A-tailing, adaptor ligation, and PCR. Libraries' quality control and quantification were performed using Qubit 3.0 Fluorometer (Life Technologies) and Agilent’s 4200 TapeStation System (Agilent), respectively. Sequencing was performed in a NextSeq2000 (Illumina) (Read 1: 28cycles, i7 Index: 10cycles, i5 Index:10 Read 2: 90cycles) at an average depth of 45,000 reads/cell followed by computational alignment using CellRanger (version (v) 6.1.1, 10x Genomics).

Single-cell RNA sequencing analysis

The scRNA-seq analysis of the BM samples was performed using R v 4.1.3 and Seurat v 4.3.0^5^. For data preprocessing, ambient RNA detection was performed using SoupX v.1.6.2^6^, and doublet scores were calculated by scDblFinder v.1.8.0^7^. Datasets were filtered individually based on a library complexity of more than 200 features, features detected in more than three cells, doublets, and high percentages of mitochondrial genes (>5%). Because the exploratory data analysis revealed potential ambient RNA contamination, immunoglobulin genes were excluded from the downstream analysis of niche cells.

For the transcriptomic analysis during disease progression, the Control, MGUS and MM cell datasets were merged, normalized using the SCTransform method^8^, and analyzed by principal-component analysis (PCA) on the most variable genes (k = 2,000) across all cells. With the principal components (PCs) (k = 30), unsupervised clustering was performed by computing the K-nearest neighbors, applying the Louvain algorithm at resolution 0.4, and cells were projected in two dimensions using Uniform Manifold Approximation and Projection (UMAP)^9^. Significantly upregulated genes in each cluster compared to all other clusters (Bonferroni-adjusted p-values <0.05) were identified using the Seurat Function FindMarkers. Manual cell type annotations identified cell types according to published canonical marker genes. The same analytical approach was applied to analyze the VRd treatment dataset individually.

To characterize BM cells at a high resolution, endothelial cells (EC), and mesenchymal cells (MSC) were subsetted into separate objects. Additionally, outlier cells within EC and MSC, lacking canonical marker genes for their respective population, were excluded from downstream analysis.

***Differential Expression analysis***

Differentially expressed genes (DEGs) were determined with the Seurat FindMarkers function for transcripts detected in at least 30% of cells using a log_2_-fold-change (log2FC) threshold of 0.5 with MAST (Model-based Analysis of Single-cell Transcriptomics) methodology^10^. An adjusted P-value threshold of 0.05 based on Bonferroni multiple testing correction was used as a cutoff to select significant DEGs. Significant DEGs for each group were used as input for a Gene Ontology (GO) Over-representation analysis (ORA) using the clusterProfiler R package^11^ to investigate the biological differences between groups, GO terms with corrected P-values less than 0.05 were considered significantly enriched. Gene Set Enrichment Analysis (GSEA) was also calculated using the GO and Hallmark gene sets from the Molecular Signatures Database v.7.5.1 in the list of DEGs ranked according to log2FC. Heatmap plots were generated using the SCP package v.0.5.4^12^.

***Signature score***

The MM interferon (IFN)-signature score was computed using Seurat’s AddModuleScore function using the top 50 DEGs of the MM2 EC subgroup calculated with the Seurat FindMarkers function (parameters min.pct=0.4 thresh.use=1, test=’MAST’). This signature score was applied across all cell types, including the treatment dataset, to assess IFN pathway activation and. The resulting score provides a quantitative measure that reflects how strongly a particular gene expression profile—often derived from a set of genes associated with a biological process or cell state—is expressed in a given cell or population. Unlike MM2 EC, which formed a clearly separated cluster based on their transcriptomic profile, MM MSC with elevated IFN signaling were defined based on a signature score exceeding a threshold of 0.25.

Sex inference

To ensure that the identified cluster was not biased by unequal contributions from individual mice—particularly since both sexes were included in the pooled samples—we inferred the sex of origin for each cell using two complementary approaches (**Supplementary Fig. 3**). First, PCA was computed using (i) the expression of sex-specific genes and (ii) exclusively Y chromosome–encoded genes. Both analyses revealed separation of cells by sex as denoted by the segregation of cells with more than one count for Y-linked genes. Second, we calculated the expression ratio of Y-linked genes (*Uty, Eif2s3y*) to X-linked genes (*Tsix, Xist*) to assign the most likely sex of origin to each cell following the strategy previously described^13^. Together, these analyses confirmed that both male- and female-derived cells contributed to all EC and MSC states, supporting that the observed cluster represent pooled contributions from multiple animals rather than being dominated by any single individual or sex.

Gene Regulatory Network analysis

Gene Regulatory Network (GRN) activity was interrogated following the workflow implemented by SCENIC^14^. This approach identifies regulons—sets of genes regulated by the same transcription factor (TF), acting like coordinated genetic programs. An equal number of cells per condition and cell type was randomly selected for the analysis. Briefly, all the genes were trained in the GENIE3 package and used to develop a random forest model for selecting co-expression modules between TFs and target genes (regulons). Regions for TFs searching were restricted to a 10-k distance centered on the transcriptional start site (TSS) or 500 bp upstream of the TSSs. Then, RcisTarget was used to refine the regulons by inferring direct targets of the transcription factors. False-positive were indirectly targeted by TFs binding motifs, and candidate TFs were removed. Finally, regulon activity scores were calculated for each cell, using the AUCell package, to determine whether the regulons were in an active or inactive state. Specifically, to identify the master regulators of "MM2" IFN EC, regulon analysis was first computed by comparing "MM2" EC to non-IFN MM EC and then extended to include all ECs across disease stages (**Figure 2D; supplemental Figure 4A-C**).

Cell-to-cell communication analysis

To assess receptor-ligand interactions between the two BM niche populations (EC and MSC) and the myeloma PC, we used Liana v.0.1.13^15^ with five different methods for predicting cell-cell interactions (cellphonedb, connectome, NATMI, logFC, and SingleCellSignalR). The results only retain interactions specifically between one cell type and another while excluding interactions within each cell type. The liana_aggregate function was used to generate an aggregate rank score for each interaction, reflecting only the specificity of interactions, and interactions were filtered based on their statistical significance (p-value < 0.01). Further bootstrapping analysis was conducted, and an equal number of cells were selected from each cell type. Subsequently, the LIANA algorithm was employed to predict interactions, iterating this process 100 times for robustness. The outcomes were assessed based on the percentage of predictions for each interaction across these iterations. The OmnipathR R package^16^ was used to import functional annotations for all genes involved in ligand-receptor interactions. To illustrate the strength of specific interactions between different groups, several ligand-receptor pairs with high mean values were selected for visualization.

**Immunofluorescence staining of mice BM samples**

Femurs were collected from control (YFP_cγ1_), MGUS (BI_cγ1_), MM (BI_cγ1_ and MI_cγ1_), alongside VRd-treated humanized BI_cγ1_-Crbn^I391V^ mice after 20 weeks of therapy. Bone tissue sections were fixed in formol 4% (PanReac) for 24 hours at RT and then decalcified using EDTA 0.25 M pH 6.95 (Invitrogen) for 10 days at RT with agitation. Following decalcification, the samples were washed in distilled water for 5 min and sequentially dehydrated in ethanol at increasing concentrations: 70% for 1 hour, 80% for 1 hour, 96% for 1 hour, and 100% overnight. The samples were then cleared in xylol (PanReac) for 4 hours. Fixed bone samples were subsequently embedded in paraffin and incubated at 60 °C overnight. Subsequently, 4 µm tissue sections were mounted on microscopy slides and dried in a desiccator at 37 °C overnight. The preparations were deparaffinized and rehydrated through a series of graded alcohols. Antigen retrieval was performed by heating the slides in 10 µM Citrate (pH6) for 30 minutes at 95°C.

For immunofluorescence (IF) staining, the Formalin-Fixed Paraffin-Embedded (FFPE) sections were blocked with Avidin/Biotin Blocking Kit (Abcam) following manufacturer indications and, subsequently, with BSA 5% for 30 min at RT. The following primary antibodies were used: rat anti-Endomucin antibody (V.7C7) (Santa Cruz Technology 1: 50 dilution), goat Leptin R Biotinylated Antibody (R&D Systems, part of Bio-Techne, 1: 50 dilution), rabbit GBP2 Polyclonal Antibody (Proteintech, 1: 100 dilution) and rabbit ISG15 Polyclonal Antibody (Proteintech, 1: 100 dilution).

The slides were subsequently incubated with the corresponding secondary antibodies in a 1:200 dilution: Goat anti-Rabbit IgG (H+L) Highly Cross-Adsorbed Secondary Antibody, Alexa Fluor™ 647 (Invitrogen), Goat anti-Rabbit IgG (H+L) Highly Cross-Adsorbed Secondary Antibody, Alexa Fluor™ 488 (Invitrogen), Goat anti-Rat Alexa Fluor™ 568 (Invitrogen) and Streptavidin, Alexa Fluor™ 488 Conjugate (Invitrogen). Nuclei were stained with DAPI (Vectashield 1:50 dilution). Fluorescence images were acquired using the Vectra Polaris Multispectral Imaging System (Perkin Elmer).

Human

For human study, clinical BM aspirate samples from individuals of both sexes with newly diagnosed MGUS (n = 5), smoldering multiple myeloma (SMM) (n =2), or MM (n =10) were obtained from the University of Navarra Biobank. Human BM samples from healthy adult donors (50-80 years of age), healthy donors (HD) (n=8) undergoing orthopedic surgery (hip or knee replacement) from Hospital Universitario de Navarra (HUN). For IF staining, archived paraffined human BM biopsies of MGUS (n=4) and MM (n=7) patients were obtained from the Pathology Department of Clínica Universidad de Navarra (CUN). Bone chips from HD (n=2) undergoing orthopedic surgery (hip or knee replacement) from HUN were also used as control. The clinical characteristics of all individuals are described in **supplemental Table 2**. All samples were collected, processed, banked, and thawed for further processing. This study was approved by the University of Navarra's Institutional Review Board (Project 2022.100mod1) and the Navarra Department of Health Ethics Committee for Drug Research (CEIm, PI_2022/92 MS-1) and conducted under the Declaration of Helsinki. Informed consent was obtained from all patients. Personal data was kept confidential following the Organic Law 3/2018 on personal data protection and Spanish Law 14/2007 on Biomedical research. All collection samples are codified; only authorized personnel can correlate the patient’s identity with the codes.

Isolation and fluorescence-activated cell sorting of human BM mesenchymal-osteolineage primed cells

After sample collection, red blood cells were lysed twice for 15 minutes at room temperature with rotation following the ratio of 45 ml of ACK lysis buffer per 5 ml of human sample. In the case of samples from healthy adult donors, portions of bones present in the samples were cut into small pieces, and after being crushed, the remaining fragments were collagenized with the appropriate volume of PBS (3ml/sample) with 0.3% collagenase I and dispase (5U/ml) for 15 min at 37ºC and shaking at 200 rpm. FBS, representing 10% of the digestion volume, was added to stop the digestion of collagenase. After digestion, the collagenized fractions were filtered through a 70 μm filter into a collection tube and pooled into one sample. To determine live cell concentration and viability, 10 μl of each sample was stained with acridine orange and propidium iodide (AO/PI) solution (Nexcelom) and analyzed with Cellometer K2 Image Cytometer (Nexcelom Bioscience). Cells were subsequently cryopreserved with FBS and 10% dimethyl sulfoxide (DMSO, Sigma-Aldrich) in liquid nitrogen until use.

Frozen BM samples were thawed at 37ºC in dextran-40 solution supplemented with 25% human albumin at 20%. Cell viability of thawed cells was assessed using Nexcelom Cellometer as described above. The sample was then centrifuged and stained for 30 min on ice with the following combination of conjugated antibodies at a concentration of 1/100 except anti-Lin (3 μl/test- test 25x10^6^cells): BV510 labeled anti-Lin (including CD3, CD10, CD19, CD45, and CD64), BV421 labeled anti-CD235, BV421 labeled anti-CD45, FITC labeled anti-CD31, APC-Cy7 labeled anti-CD9, PE labelled anti-CD146, and PerCP-Cy5.5 labelled anti-CD271. First, dead cells and debris were excluded by FSC, SSC and adding 10 μl of TO-PRO-3. BM niche MSC were prospectively isolated based on the following immunophenotype: TO-PRO-3^-^/Lin^-^/CD45^-^/CD235^-^/CD16^-^/CD56^-^/CD31^-^/CD271^+^/CD146^+/-^. For bulk RNA-sequencing (bulk RNA-seq) studies, MSC were sorted in 100 μl of Lysis/Binding Buffer (Ambion), vortexed, and stored at -80ºC until further processing.

Bulk RNA sequencing

Total RNA of human BM MSC sorted from old individuals and patients were isolated with the MagMAX mirVana Total RNA Isolation Kit (Applied Biosystems, Thermo Fisher) according to the manufacturer’s protocol but finally resuspended in 15ul of ultrapure water and examined using Takara’s SMART-Seq v4 plus kit (Cat. No. R400753, Takara Bio.) as indicated by the manufacturer’s instructions. Briefly, oligo(dt) primers are used to obtain a full-length cDNA through the SMART® (Switching Mechanism at 5′ end of RNA Template) technology. Once full-length cDNA is synthesized, libraries are prepared through an enzymatic fragmentation method, followed by library amplification and indexing sequencing libraries using unique dual indexes (Cat. R400745, Takara Bio). Libraries were quantified with Qubit dsDNA HS Assay Kit, and their profile was examined using Agilent’s HS D5000 ScreenTape Assay. Sequencing was carried out in an Illumina NextSeq2000 using paired-end dual-index sequencing (Rd1: 60 cycles; i7: 8 cycles; i5: 8 cycles; Rd2: 60 cycles) at a depth of 10-20 million reads per sample.

Raw reads were demultiplexed via bclfastq2 Conversion Software v 2.20 (Illumina) to convert base call (BCL) files into FastQ files. Quality control of FastQ files was performed using fastQC (Bioinformatics Babraham Institute), and reads were trimmed for contaminating sequence adapters and poor-quality bases using Trimmomatic^17^. Sequencing reads were aligned to hg19 human reference using STAR v 2.6^18^. Uniquely aligned reads were assigned to GENCODE18 gene annotations using HTSeq v 0.11^19^ and quantified with htseq-counts run in stranded mode with default parameters.

Bulk RNA sequencing analysis

Before proceeding with the analysis, we extensively evaluated the MSC proportion in our samples. First, we used CIBERSORTx deconvolution platform to estimate the relative cell fractions of MSC and contaminating cells present in our previous work with human scRNA-seq experiments (**supplemental** **Figure 16A**). The signature matrix was prepared using pseudobulks of EC, MSC, neutrophils, megakaryocytes, T cells, NK, dendritic cells, PC, and B cells. Gene counts were normalized for MSC (excluding the pseudobulk for other populations) using the voom-limma R package after removing genes with low mean expression (less than 10 counts). The signature matrix from our monoclonal gammopathies samples was uploaded to the CIBERSORTx website according to its instructions. Permutations were set to 500, and the rest of the parameters retained the default. After running CIBERSORTx, we obtained the relative proportions of MSC (including pericytes) and the contaminating populations in each sample. Next, gene signature enrichment analysis using the AUCell^14^ method was applied to quantify each sample´s mesenchymal signatures. First, we used the Gene Set function from the GSEAbase R package to create a gene list of 20 markers that define the MSC. After processing the gene sets, the genes were ranked for each sample from the ones with the highest expression to those with the lowest using the AUCell´s AUC_buildRanking function.

After determining the proportions of MSC in our samples and confirming the identity by the expression of MSC canonical markers (**supplemental** **Figure 16B**), we proceeded with the subsequent analysis. We filtered those genes with an average expression per condition lower than 2 CPM (counts per million). Next, the batch effect was corrected using the sva package (v 3.46)^20^. These gene count estimates were normalized, prefiltered, and used for determining the differential expression of genes between the different groups through the DESeq2 package^21^. In this differential expression analysis, the proportions of MSC estimated by CIBERSORTx were included in the model as a covariate to avoid its possible impact on the results. The Benjamini-Hochberg False discovery rate was applied to adjust for multiple hypothesis testing, and genes with an adjusted p-value below 0.05 were considered differentially expressed. To evaluate the contribution of contaminating cells to differential gene expression we calculated the Spearman correlation between the expression of previously identified DEGs and the niche proportions quantified with CIBERSORTx, and the cell-type signature score calculated with AUC. Genes whose expression was significantly correlated with any of those two variables with an adjusted p-value below 0.05 were excluded from the results. Next, to deal with the high variability observed among the samples, we performed a Wilcoxon Test, where we selected those genes with an adjusted p-value below 0.05. This set of genes was the one used for the downstream analysis.

ORA using the functions enrichGO and compareCluster in the package clusterProfiler (v 4.6.2)^11^ were conducted to interpret the enrichment and pathways of the filtered DEGs. Gene sets with an adjusted p-value below 0.05 were considered statistically significant. Additionally, GSEA was calculated using the Hallmark gene sets from the Molecular Signatures Database v.7.5.1 in the list of DEGs ranked according to log2FC. In addition, the AUCell method was applied to quantify the IFN signature discovered in the mouse data.

The transcription factor activity was estimated by Decoupler^22^. It compiles information from CollecTRI^23^ database via Omnipath package^16^, and applies a Univariate Linear Model (ulm) to normalized log-transformed RNA counts so it can predict the observed gene expression based on the TF's TF-Gene interaction weights.

**Immunofluorescence staining of human BM biopsies**

The bone chips from HD were processed for IF staining following the protocol previously described for mouse samples. Subsequently, FFPE samples from HD and archived human BM biopsies were blocked with 5% BSA for 30 minutes at room temperature (RT) before incubation with the following primary antibodies: mouse anti-CD271 monoclonal antibody (NGF Receptor) (ME20.4) (Invitrogen™ 1: 50 dilution), mouse anti-CD31/PECAM-1 antibody (Biotechne 1: 40 dilution), rabbit GBP2 Polyclonal Antibody (Proteintech, 1: 100 dilution) and rabbit ISG15 Polyclonal Antibody (Proteintech, 1: 100 dilution).

The slides were subsequently incubated with the corresponding secondary antibodies in a 1:200 dilution: Goat anti-Rabbit IgG (H+L) Highly Cross-Adsorbed Secondary Antibody, Alexa Fluor™ 488 (Invitrogen), goat anti-Mouse IgG (H+L) Cross-Adsorbed Secondary Antibody, Alexa Fluor™ 568 (Invitrogen). Nuclei were stained with DAPI (Vectashield 1:10 dilution). Fluorescence images were acquired using the Vectra Polaris Multispectral Imaging System (Perkin Elmer).

**Immunofluorescence analysis of human BM biopsies**

This analysis assessed mesenchymal IFN-enriched cells by evaluating the coexpression of MSC marker and IFN-responsive genes. The percentage of MSC was quantified based on CD271expression, while IFN-enriched cells were identified through staining for IFN-responsive genes, including GBP2 and ISG15. Image analysis and quantification were conducted using QuPath (version 0.5.1), providing a detailed assessment of IFN-driven mesenchymal alterations.

Statistical analyses

Statistical analysis was carried out in R (v 4.1.3). All results in the graphs are means ± Standard error of the mean. Tests used to evaluate statistical significance are detailed in each method section.

Data availability

All the murine scRNA-seq data and raw human bulk RNA-seq data generated in this study will be publicly available as of the date of peer-reviewed publication and is available under request; contact. The public scRNA-seq datasets of the non-hematopoietic BME of newly diagnosed individuals with MM and individuals without myeloma are available on ArrayExpress, no. E-MTAB-9139, and were used to validate the enrichment of the IFN signature in a subset of MM patients^24^. The bulk RNA-sequencing datasets of MSC from individuals with MM at diagnosis and after induction treatment are available on ArrayExpress, no. E-MTAB-9285, and were used to confirm suppression of the IFN signature following therapy^24^.

**REFERENCES SUPPLEMENTAL METHODS**

1	Larrayoz M, Garcia-Barchino MJ, Celay J, Etxebeste A, Jimenez M, Perez C *et al.* Preclinical models for prediction of immunotherapy outcomes and immune evasion mechanisms in genetically heterogeneous multiple myeloma. *Nat Med* 2023; **29**: 632–645.

2	McCaughan GJ, Gandolfi S, Moore JJ, Richardson PG. Lenalidomide, bortezomib and dexamethasone induction therapy for the treatment of newly diagnosed multiple myeloma: a practical review. *Br J Haematol* 2022; **199**: 190–204.

3	Krönke J, Fink EC, Hollenbach PW, MacBeth KJ, Hurst SN, Udeshi ND *et al.* Lenalidomide induces ubiquitination and degradation of CK1α in del(5q) MDS. *Nature 2015 523:7559* 2015; **523**: 183–188.

4	Fink EC, McConkey M, Adams DN, Haldar SD, Kennedy JA, Guirguis AA *et al.* Crbn I391V is sufficient to confer in vivo sensitivity to thalidomide and its derivatives in mice. *Blood* 2018; **132**: 1535–1544.

5	Stuart T, Butler A, Hoffman P, Hafemeister C, Papalexi E, Mauck WM *et al.* Comprehensive Integration of Single-Cell Data. *Cell* 2019; **177**: 1888-1902.e21.

6	Young MD, Behjati S. SoupX removes ambient RNA contamination from droplet-based single-cell RNA sequencing data. *Gigascience* 2020; **9**: 1–10.

7	Germain PL, Lun A, Macnair W, Robinson MD. Doublet identification in single-cell sequencing data using scDblFinder. *F1000Research 2021 10:979* 2021; **10**: 979.

8	Hafemeister C, Satija R. Normalization and variance stabilization of single-cell RNA-seq data using regularized negative binomial regression. *Genome Biol* 2019; **20**: 1–15.

9	McInnes L, Healy J, Saul N, Großberger L. UMAP: Uniform Manifold Approximation and Projection. *J Open Source Softw* 2018; **3**: 861.

10	Finak G, McDavid A, Yajima M, Deng J, Gersuk V, Shalek AK *et al.* MAST: A flexible statistical framework for assessing transcriptional changes and characterizing heterogeneity in single-cell RNA sequencing data. *Genome Biol* 2015; **16**: 1–13.

11	Yu G, Wang LG, Han Y, He QY. ClusterProfiler: an R package for comparing biological themes among gene clusters. *OMICS A J Integr Biol* 2012; **16**: 284–287.

12	Hao Z. SCP: Single Cell Pipeline. 2023.https://github.com/zhanghao-njmu/SCP.

13	Gaertner Z, Oram C, Schneeweis A, Schonfeld E, Bolduc C, Chen C *et al.* Molecular and spatial transcriptomic classification of midbrain dopamine neurons and their alterations in a LRRK2G2019S model of Parkinson’s disease. *Elife* 2025; **13**. doi:10.7554/ELIFE.101035.

14	Aibar S, González-Blas CB, Moerman T, Huynh-Thu VA, Imrichova H, Hulselmans G *et al.* SCENIC: Single-cell regulatory network inference and clustering. *Nat Methods* 2017; **14**: 1083.

15	Dimitrov D, Türei D, Garrido-Rodriguez M, Burmedi PL, Nagai JS, Boys C *et al.* Comparison of methods and resources for cell-cell communication inference from single-cell RNA-Seq data. *Nature Communications 2022 13:1* 2022; **13**: 1–13.

16	Türei D, Valdeolivas A, Gul L, Palacio‐Escat N, Klein M, Ivanova O *et al.* Integrated intra‐ and intercellular signaling knowledge for multicellular omics analysis. *Mol Syst Biol* 2021; **17**: 9923.

17	Bolger AM, Lohse M, Usadel B. Genome analysis Trimmomatic: a flexible trimmer for Illumina sequence data. 2014; **30**: 2114–2120.

18	Dobin A, Davis CA, Schlesinger F, Drenkow J, Zaleski C, Jha S *et al.* Sequence analysis STAR: ultrafast universal RNA-seq aligner. 2013; **29**: 15–21.

19	Anders S, Pyl PT, Huber W. Genome analysis HTSeq-a Python framework to work with high-throughput sequencing data. 2015; **31**: 166–169.

20	Leek JT, Johnson WE, Parker HS, Jaffe AE, Storey JD. The sva package for removing batch effects and other unwanted variation in high-throughput experiments. *Bioinformatics* 2012; **28**: 882.

21	Love MI, Huber W, Anders S. Moderated estimation of fold change and dispersion for RNA-seq data with DESeq2. *Genome Biol* 2014; **15**: 1–21.

22	Badia-I-Mompel P, Vélez Santiago J, Braunger J, Geiss C, Dimitrov D, Müller-Dott S *et al.* decoupleR: ensemble of computational methods to infer biological activities from omics data. *Bioinformatics advances* 2022; **2**. doi:10.1093/BIOADV/VBAC016.

23	Müller-Dott S, Tsirvouli E, Vazquez M, Ramirez Flores RO, Badia-I-Mompel P, Fallegger R *et al.* Expanding the coverage of regulons from high-confidence prior knowledge for accurate estimation of transcription factor activities. *Nucleic Acids Res* 2023; **51**: 10934–10949.

24	de Jong MME, Kellermayer Z, Papazian N, Tahri S, Hofste op Bruinink D, Hoogenboezem R *et al.* The multiple myeloma microenvironment is defined by an inflammatory stromal cell landscape. *Nature Immunology 2021 22:6* 2021; **22**: 769–780.

SUPPLEMENTAL FIGURES

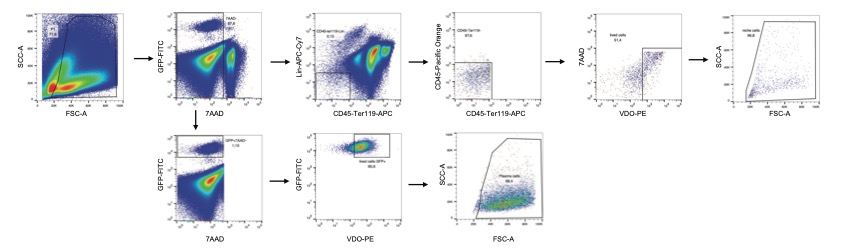
 **Supplemental Figure 1: Cell sorting strategy for murine scRNA-seq of BM cells**. Cell sorting strategy for the isolation of mouse BME cells (Lived non-hematopoietic BM cells: 7-AAD^-^ Lin^-^ CD45^-^ Ter119^-^ VDO^+^) and PC (Lived PC: 7-AAD^-^ VDO^+^ GFP^+^).

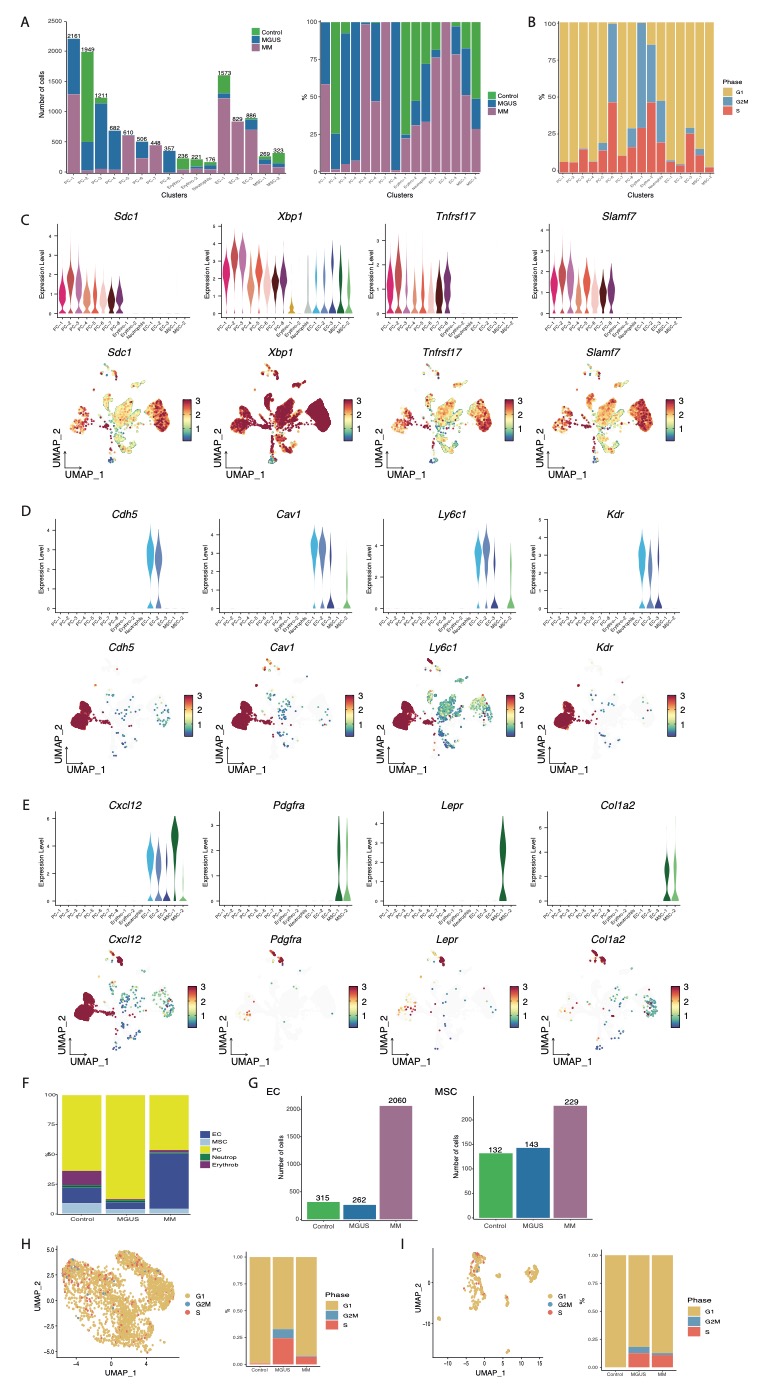

**Supplemental Figure 2: scRNA-seq analysis of murine BM cells**. (A) Left panel: bar plots showing the number of cells per cluster and dataset; Right panel: stacked bar plot representing the proportion of cells per cluster and dataset. (B) Stacked bar plot of the proportion of cells per cycle stage in each cluster. (C) Upper panel: Violin plot of gene expression of well-known markers for PC. Bottom panel: UMAP visualization of representative canonical markers for PC. (D-E) Similar to C for EC (D) and MSC (E). (F) Stacked bar plot of the proportion of BM cell populations per disease dataset. (G) Bar plot depicting the total number of cells per stage for EC (left) and MSC (right) after final filtering (see methods). (H) Analysis of cell cycle stage for EC. (I) Analysis of cell cycle stage for MSC.

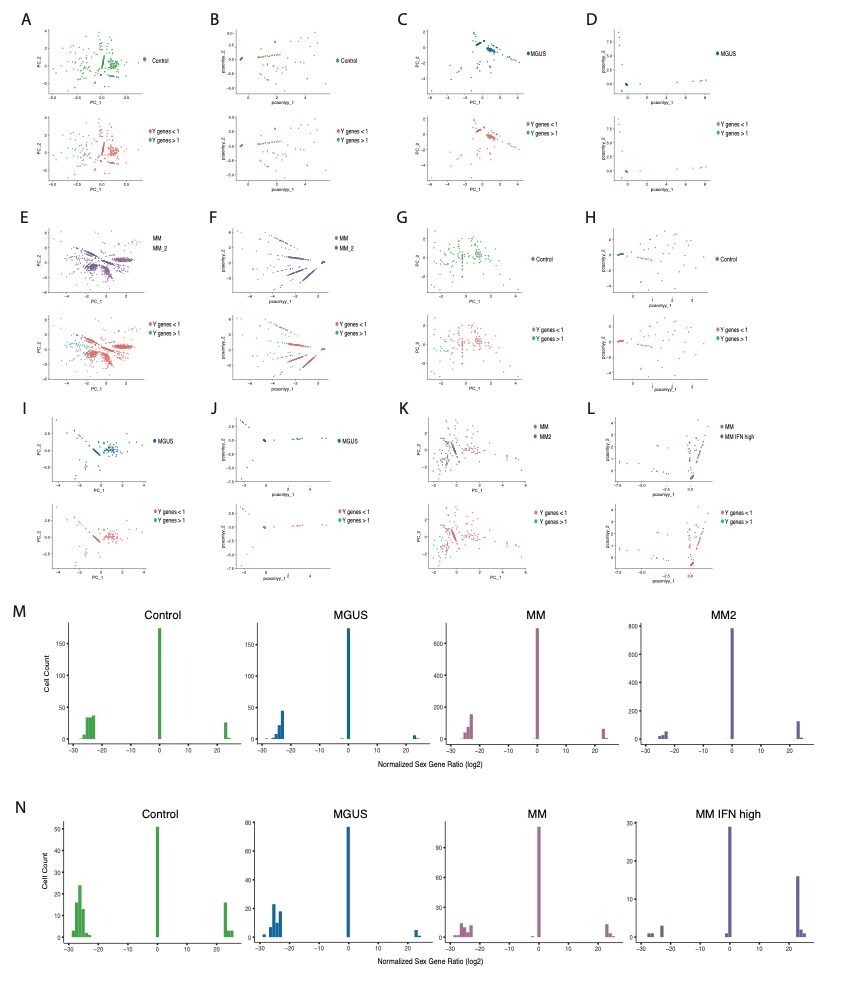

**Supplemental Figure 3: Analysis of mouse contribution to control, MGUS, and MM BI_cγ1_ EC and MSC using sex-specific genes.** (A) PCA plot showing separation of control EC based only on the expression of sex-specific genes colored by disease (upper panel) and Y-chromosome gene expression, where blue denotes cells with more than one count in Y-chromosome genes (bottom panel). (B) PCA plot showing separation of control EC based only on the expression of Y-chromosome genes colored by disease (upper panel) and Y-chromosome gene expression, where blue denotes cells with more than one count in Y-chromosome genes (bottom panel). (C,D) Similar to A,B for MGUS EC. (E,F) Similar to A,B for MM EC. MM EC cells were split into MM and MM2 clusters. (G,H) Similar to A,B for Control MSC. (I,J) Similar to A,B for MGUS MSC. (K,L) Similar to A,B for MM MSC. MM MSC cells were split into MM IFN low and high score. (M) Histogram of number of EC plotted by male to female gene ratios (scored continuously along the x-axis) in control, MGUS, and MM BI_cγ1_ EC. MM EC cells were split into MM and MM2 clusters. Three discrete peaks emerge, representing cells likely originating from either sex or those with indeterminate ratios due to technical drop-off in scRNA-seq reads. (N) Similar to M for MSC.

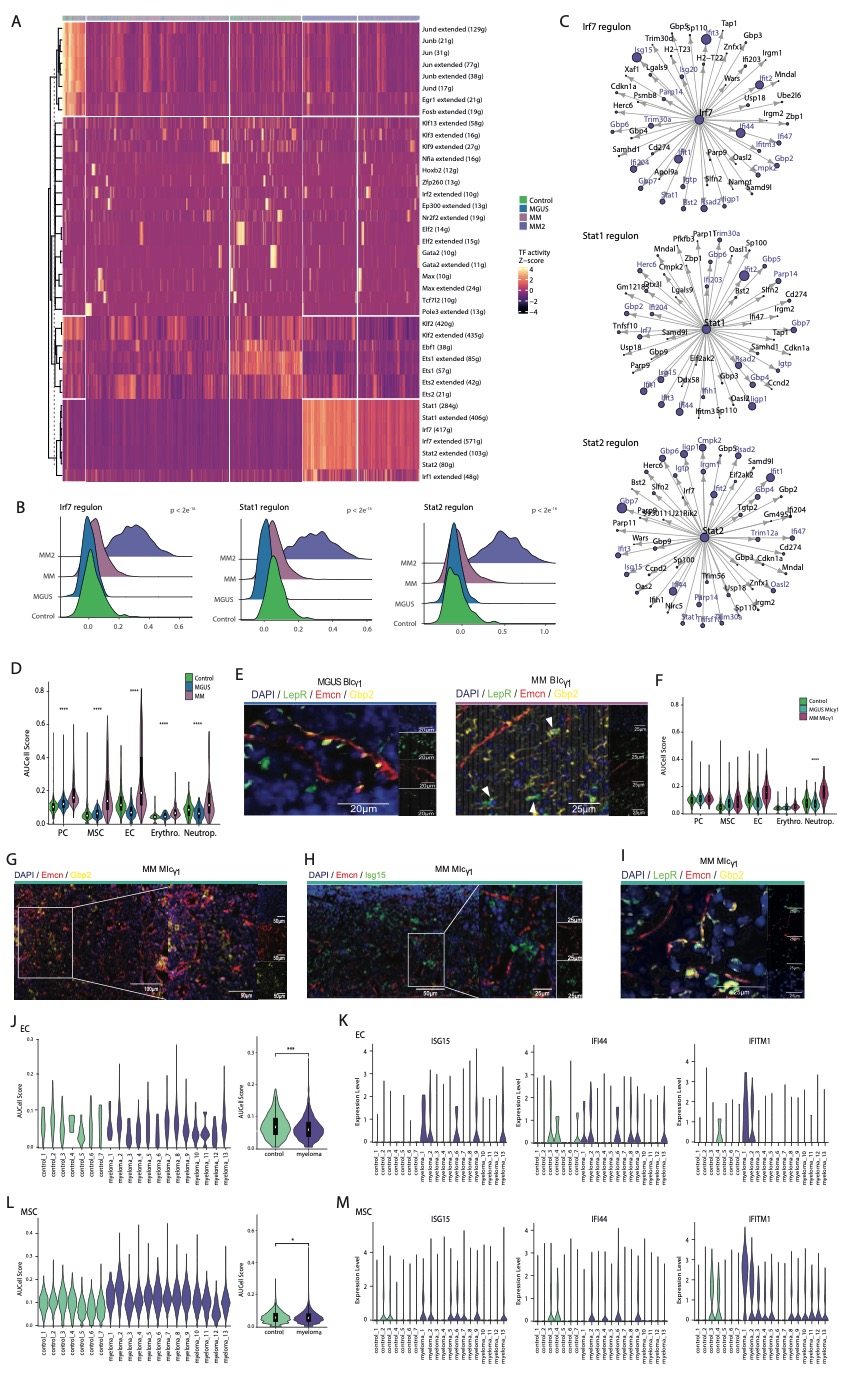
 **Supplemental Figure 4: Additional information on the MM IFN-related signature, validation in the MI_cγ1_ model and scRNA-seq human data**. (A) Heatmap showing the activities of the enriched regulons in EC (Z-score of regulon activity). (B) Density plots showing the score of Irf7, Stat1, and Stat2 regulons in EC (p. = adjusted p-value). (C) Regulon network of Irf7, Stat1, and Stat2 representing the top 50 target genes. (D) Violin plots showing AUC-score for MM IFN-related signature within BM populations in the BI_cγ1_ model. (E) IF staining of EC (Emcn, red), MSC (LepR, green), IFN response gene marker of MM2 cluster (Gbp2, yellow), and nucleus (DAPI) (blue) in FFPE femurs from MGUS BI_cγ1_ and MM BI_cγ1_ mice. White arrows point MSC LEPR+ GBP2+ (F) Violin plots showing AUC-score for MM IFN-related signature within BM populations in the MI_cγ1_ model. (G-I) IF staining of EC (Emcn, red), LEPR (green, (I)),and IFN response genes markers of MM2 cluster (Gbp2 (yellow, (G)); Isg15 (green, (H))),and nucleus (DAPI) (blue) in FFPE femurs from MM MI_cγ1_ mice. (J) Violin plots showing AUC-score for MM IFN-related signature in EC from healthy controls and MM human patients from scRNA-seq published data^24^ by individuals (left panel) and grouped by disease (right panel). (K) Violin plots comparing the distribution of the gene expression of *ISG15*, *IFI44* and *IFITM1* in EC from MM patients and healthy controls. (L-M) Similar to I-J for MSC from the same healthy controls and MM human patients from scRNA-seq published data^24^.

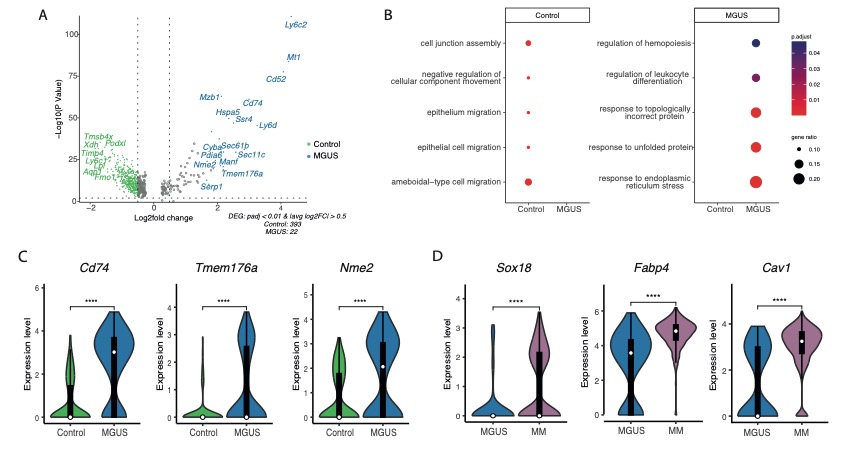
 **Supplemental Figure 5: Additional visualizations of transcriptional differences detected between EC from control, MGUS, and MM excluding MM IFN cells.** (A) Volcano plots for the differential expression analysis between EC from control and MGUS. The y-axis represents the -log_10_(p-value), and the x-axis represents the log_2_FC of the gene. The dot's color denotes the disease stage for which that DEG was detected, with grey dots representing non-significant genes. (B) GO ORA analysis between EC from control and MGUS. (C) Violin plots showing the expression of relevant upregulated genes in MGUS vs control EC. (D) Violin plots showing the expression of upregulated genes in EC from MM compared to MGUS.

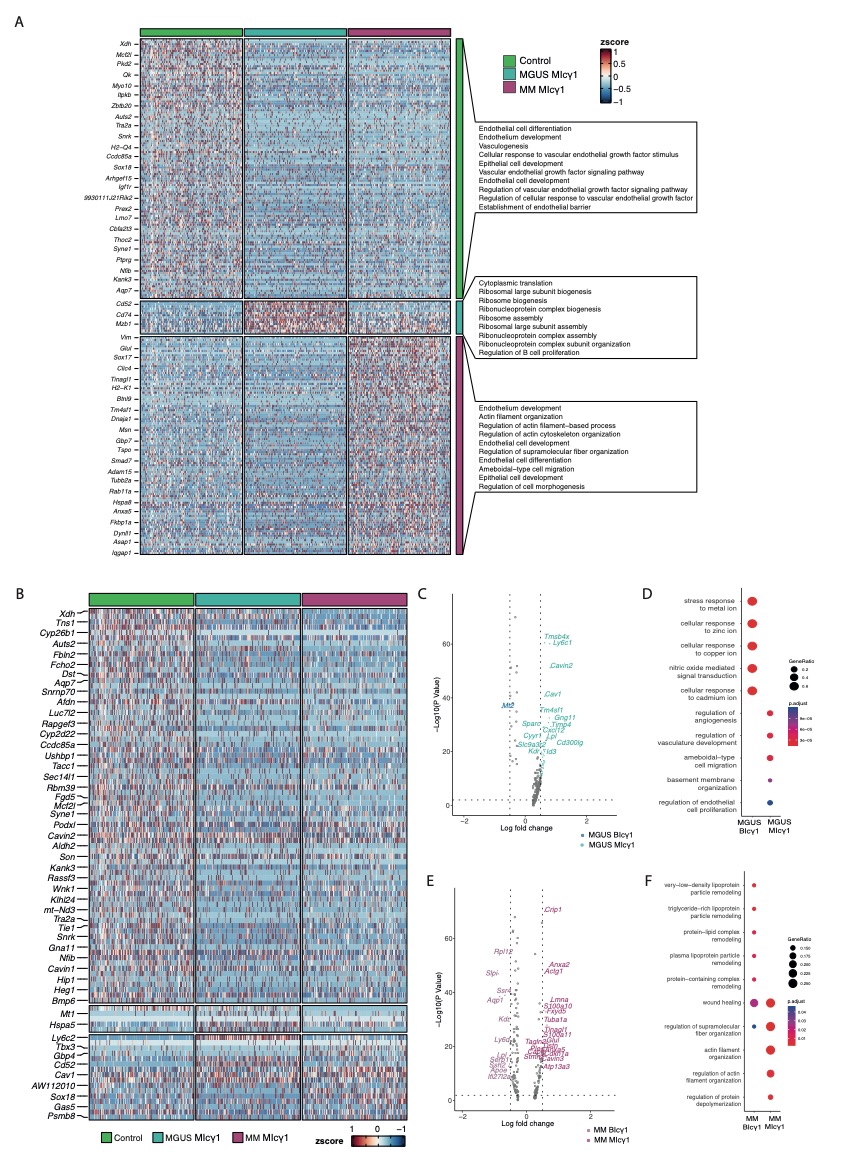
 **Supplemental Figure 6: Transcriptional changes in EC during myeloma progression in the MI_cγ1_ model.** (A) Heatmap of differential gene expression analysis in EC across control, MGUS, and MM. Enriched GO terms were identified for each disease stage, and the top 10 enriched terms are flagged on the right. Color represents the z-score scaled expression values. (B) Heatmap projecting MI_cγ1_ model gene expression of DEGs identified in EC from BI_cγ1_ at different stages of the disease. Color represents the z-score scaled expression values of EC from the MI_cγ1_ model. (C) Volcano plot of the DEGs between EC from MGUS in BI_cγ1_ and MI_cγ1_ model. The y-axis represents the -log10(p-value), and the x-axis represents the log_2_FC of the gene. The dot's color denotes the model for which that DEG was detected, with grey dots representing non-significant genes. (D) GO ORA analysis between EC from BI_cγ1_ and MI_cγ1_ model at the MGUS stage. (E-F) Similar to (C-D) for the comparison between BI_cγ1_ and MI_cγ1_ model in the MM stage.

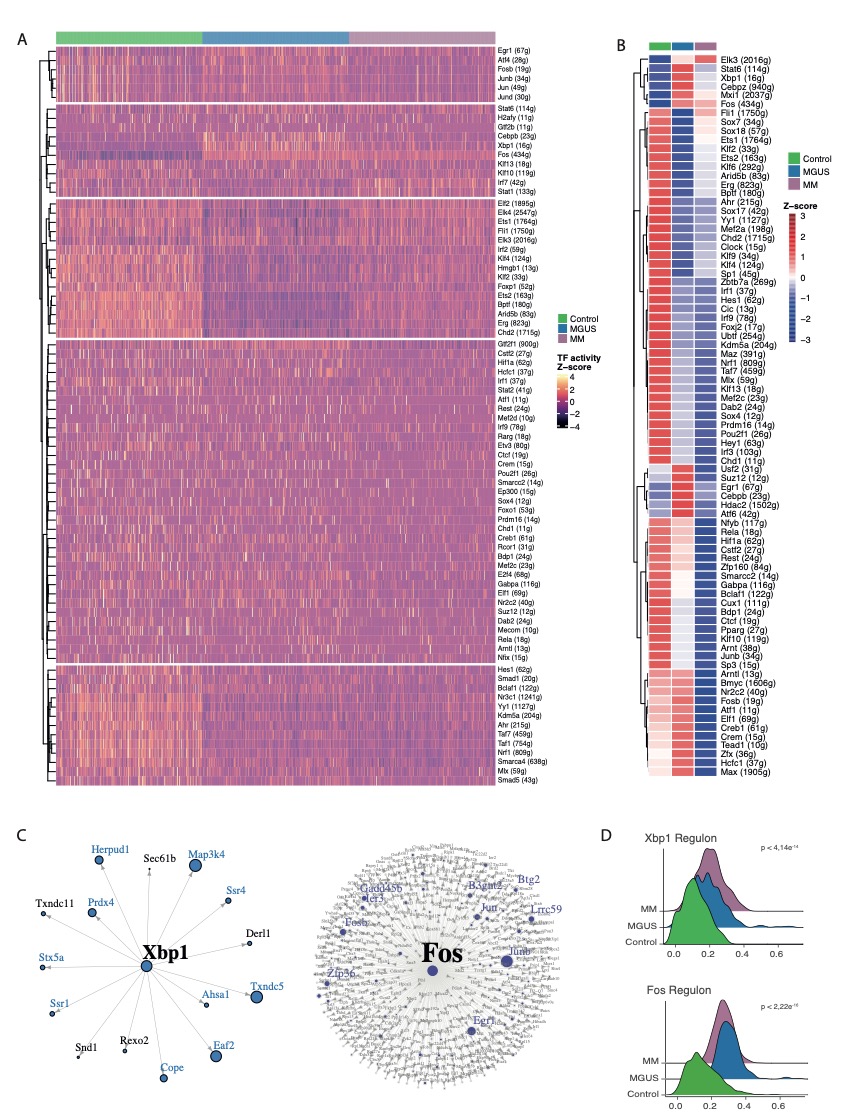

**Supplemental Figure 7: Gene regulatory network of EC during MM progression excluding MM IFN signal.** (A) Heatmap showing the activities of the enriched regulons in EC from control, MGUS and MM (Z-score of regulon activity). (B) Heatmap of mean regulon activity score of significantly enriched regulons in each stage. The color scale represents the z-score of the activity scaled by row. (C) Regulon network of Xbp1 and Fos representing the top target genes. (D) Density plots showing the score of Xbp1 and Fos regulons in EC.

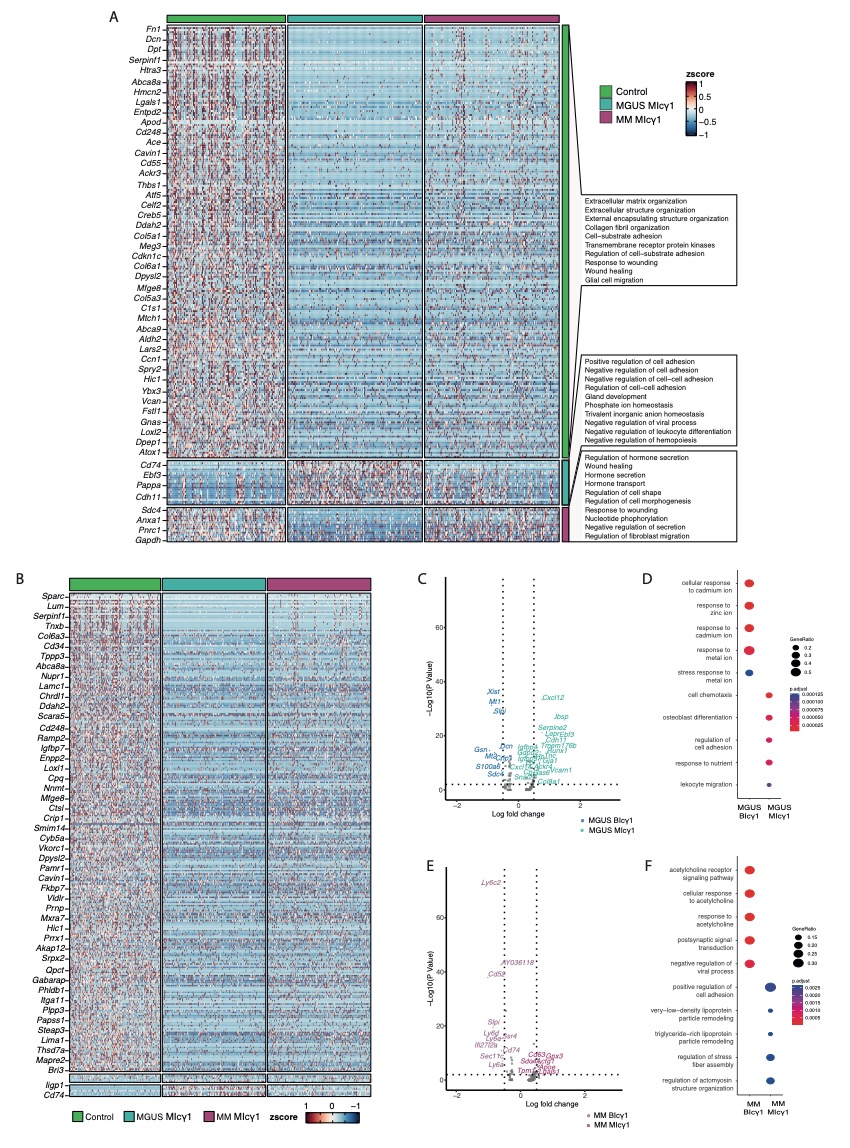
 **Supplemental Figure 8: Transcriptional remodeling of MSC during myeloma development in the MI_cγ1_ model.** (A) Heatmap of differential gene expression analysis in MSC across control, MGUS, and MM stratified by stage. Enriched GO terms were identified for each disease stage, and the top 10 enriched terms are flagged on the right. Color represents the z-score scaled expression values. (B) Heatmap projecting MI_cγ1_ model gene expression of DEG identified from BI_cγ1_ model in control, MGUS and MM. Color represents the z-score scaled expression values of MSC from the MI_cγ1_ model. (C) Volcano plot of the DEGs between MSC in MGUS from BI_cγ1_ and MI_cγ1_ model. The y-axis represents the -log10(p-value), and the x-axis represents the log_2_FC of the gene. The dot's color denotes the model from which that DEG was detected, with grey dots representing non-significant genes. (D) GO ORA between MSC from BI_cγ1_ and MI_cγ1_ model at the MGUS stage. (E-F) Similar to (C-D) for the comparison between MSC from BI_cγ1_ and MI_cγ1_ model in the MM stage.

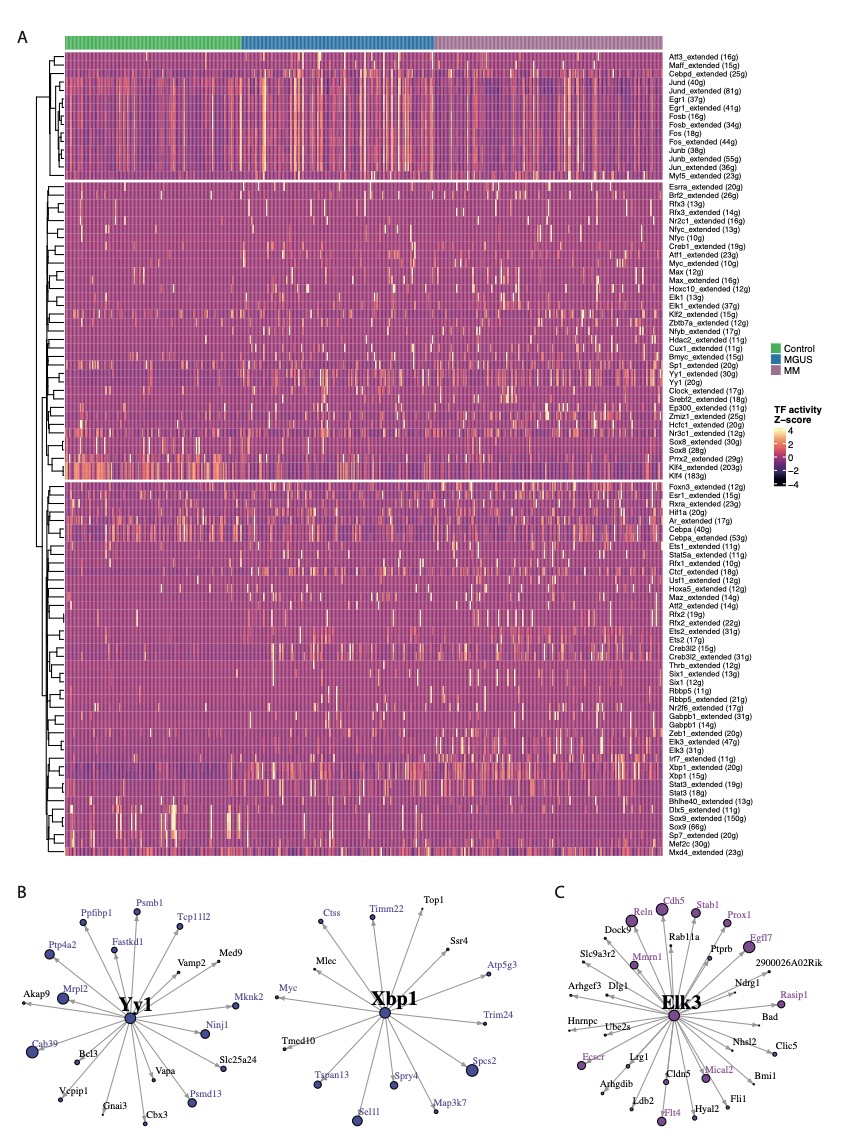
 **Supplemental Figure 9: Gene regulatory network of MSC during MM progression excluding MM IFN signal.** (A) Heatmap showing the activities of the enriched regulons in MSC from control, MGUS, and MM (Z-score of regulon activity). (B) Regulon network of Yy1 and Xbp1, representing the top associated genes (C) Regulon network of Elk3 representing the top associated genes.

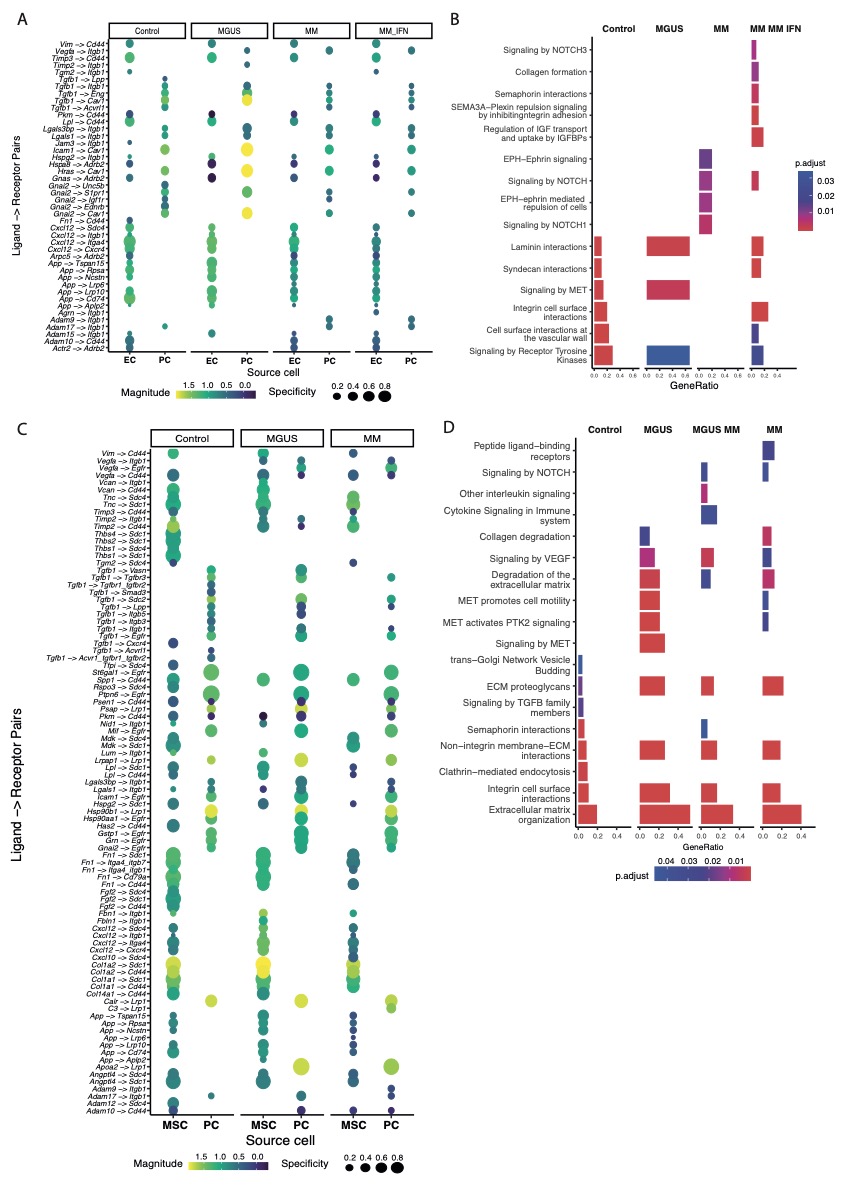
 **Supplemental Figure 10: Cell-cell communication analysis between EC, MSC and PC.** (A) Dot plots of ligand-receptor interactions between PC and EC that are maintained along the disease stages. Dot size and color indicate the specificity and the strength of the interaction, respectively. (B) Representative Reactome pathways enriched in EC-PC L-R pairs specific for each stage. (C-D) Similar to (A-B) for MSC-PC communication.

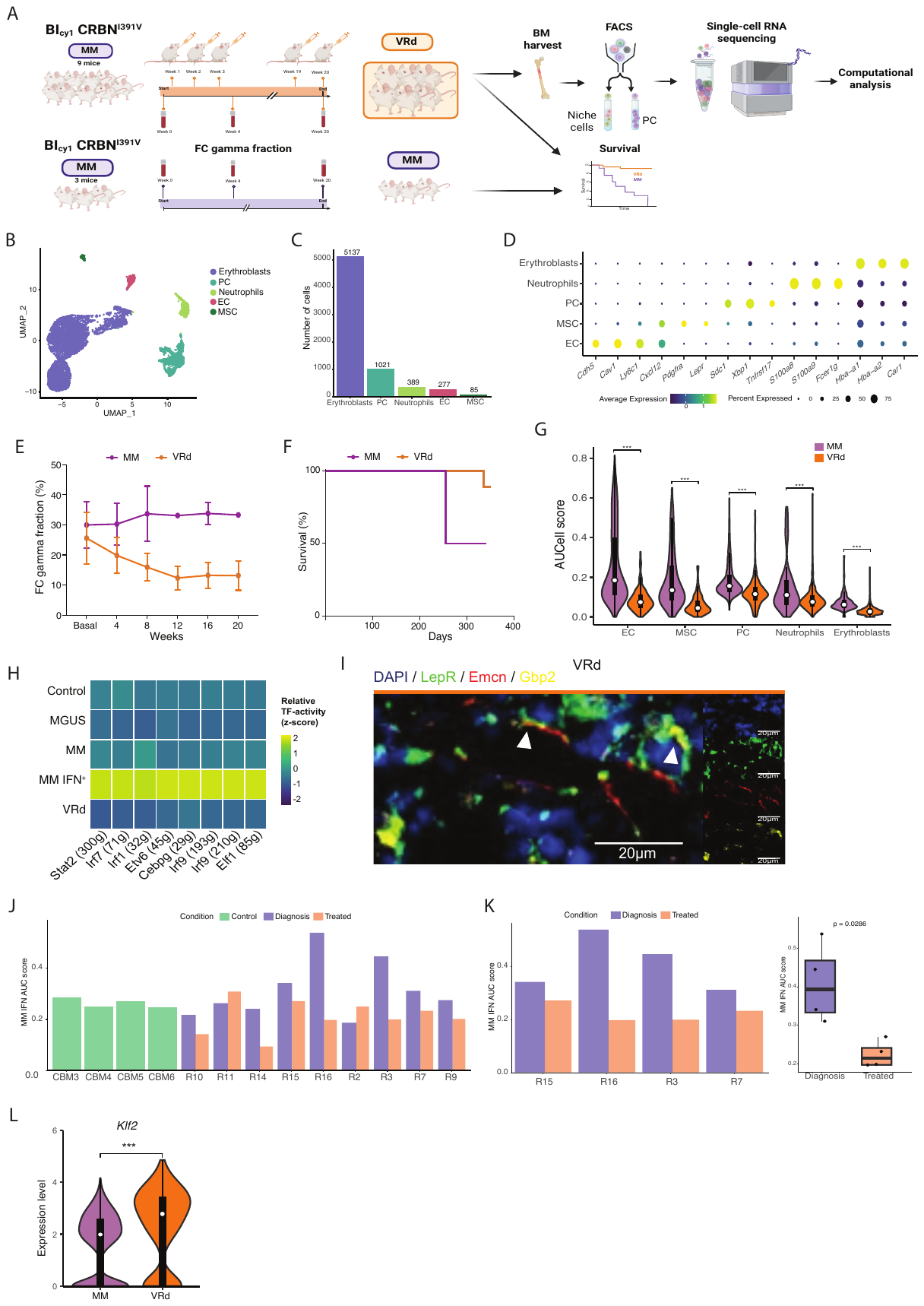
 **Supplemental Figure 11: Additional analysis of VRd-treated mice and transcriptional alterations in EC and MSC from human patients.** (A) VRd experiment overview representing treatment administration, serum collection for FC gamma fraction and scRNA-seq isolation of PC, MSC and EC from MM BI_cγ1_ *Crbn*^I391V^ (pool of six mice). MM BI_cγ1_ *Crbn*^I391V^ were used as controls for FC gamma fraction and survival. (B) UMAP representation of BM microenvironment cells from BI_cγ1_-*Crbn*^I391V^ VRd-treated mice colored by cell clustering. (C) Bar plots showing the number of cells per BM population. (D) Dot plot of canonical markers used to define PC, EC, MSC, erythroblasts, and neutrophils. The dot size reflects the percentage of cells within the cluster expressing each gene, and the color represents the average expression level. (E) Representation of the percentage of serum FC gamma fraction in VRd-treated versus MM group. (F) Kaplan–Meier survival curves of treated and non-treated MM BI_cγ1_ mice. (G) Violin plots showing AUC-score for MM IFN-related signature within BM populations in VRd treated and MM BI_cγ1_ mice. (H) Heatmap representing z-scaled mean regulon activity score of IFN-related regulons in EC across each condition. (I) IF staining of EC (Emcn, red), MSC (Lepr, green), IFN response (Gbp2, yellow), and nucleus (DAPI) (blue) in FFPE femurs from VRd-treated mice. (J) Bar plots showing the AUC-score for MM IFN-related signature in MSC from healthy controls and MM human patients before and post-induction treatment from bulk RNA-seq published data^24^ . (K) Bar plots showing the AUC-score for MM IFN-related in MSC from the subset of high IFN MM human patients before and after induction treatment^24^ by individuals (left panel) and grouped by disease (right panel). (L) Violin plot displaying the upregulation of *Klf2* in VRd-treated EC vs non-treated MM EC.

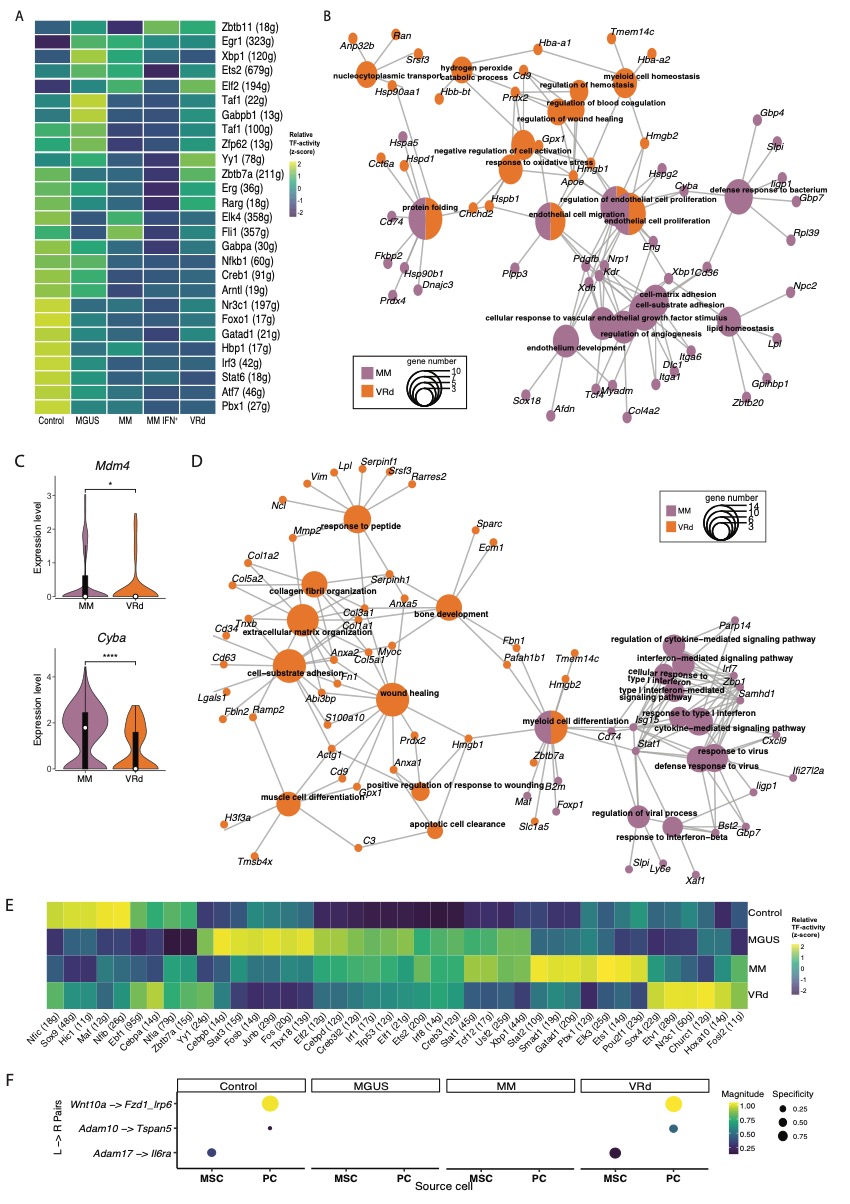

**Supplemental Figure 12: Additional information of the transcriptional remodeling of EC and MSC following VRd treatment.** (A) Heatmap representing z-scaled mean regulon activity score of inferred top specific regulons for EC in each condition. (B) Cnetplot showing the links between genes and biological processes derived from the comparison between VRd-treated and non-IFN MM EC. Node size reflects the number of significantly enriched genes in the node and colors the group condition. (C) Violin plots displaying the expression of *Mdm4* and *Cyba* in VRd-treated and MM BI_cγ1_ MSC. (D) Cnetplot showing the links between genes and biological processes derived from the comparison between VRd-treated and MM MSC. Node size reflects the number of significantly enriched genes in the node and colors the group condition (E) Heatmap representing z-scaled mean regulon activity score of inferred top specific regulons for MSC in each condition. (F) Dot plots representing L-R pairs common to control and VRd-treated mice involved in the communication between MSC and PC. Dot color represent the strength and size indicate the specificity of the interaction.

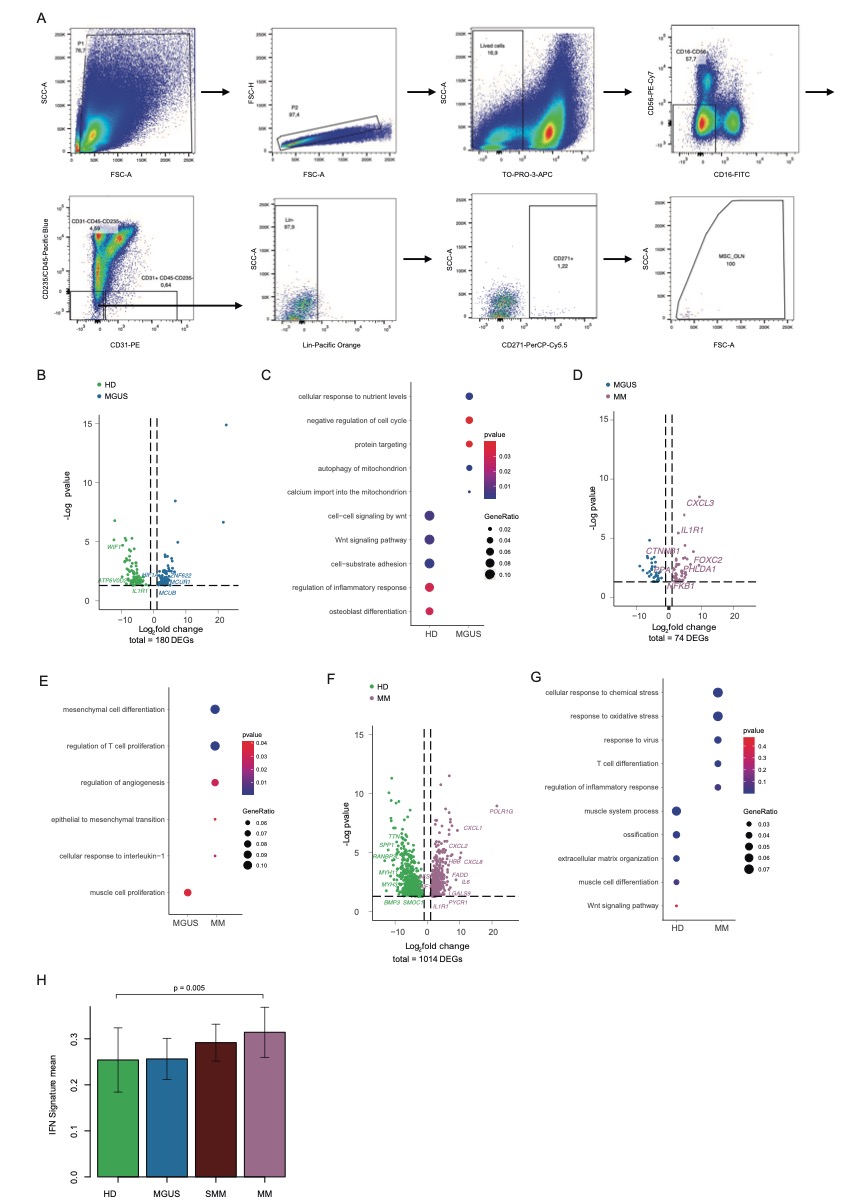

**Supplemental Figure 13: Cell sorting strategy and additional information on the transcriptional alterations of human MSC during myeloma progression.** (A) Sorting gating strategy for isolation of human BM MSC (TO-PRO-3^-^, CD16^-^, CD56^-^, CD45^-^, CD235^-^, CD31^-^, Lin^-^, CD271^+^). (B) Volcano plot of the DEGs between MGUS-MSC and HD-MSC. The y-axis represents the -log_10_(p-value), and the x-axis represents the log_2_FC of the gene. The dot's color represents the patient's disease condition. (C) GO ORA between MSC from HD and MGUS patients. (D-E) Similar to (B-C) for the comparison between MSC from MGUS and MM patients. (F-G) Similar to (B-C) for the comparison between HD-MSC and MM-MSC. (H) Bar plot depicting the mean IFN signature score across different disease conditions.

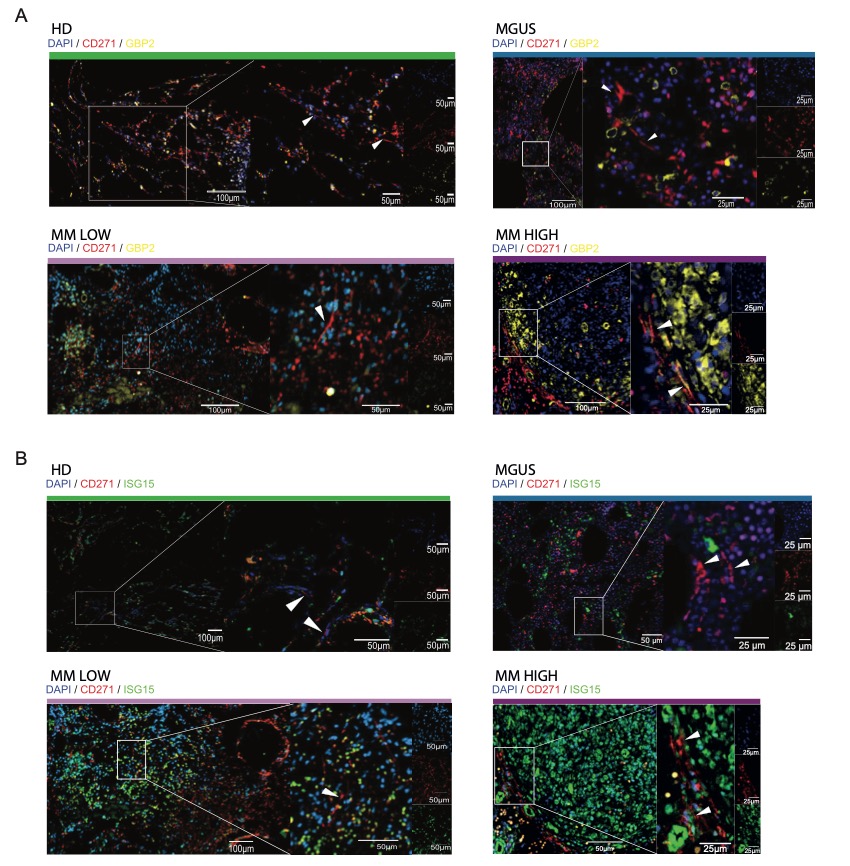
 **Supplemental Figure 14: Expanded immunofluorescence analysis of human MSC during myeloma progression.** (A-B) IF staining of MSC (CD271, red), IFN response genes (*GBP2*, yellow (A); *ISG15,* green (B)), and nucleus (DAPI) (blue) in FFPE biopsies from HD and archived paraffined human BM biopsies of MGUS and MM patients with high (MM HIGH) and low IFN signal (MM LOW) in stromal cells (see Supplementary Table 2). White arrows point MSC CD271+ in HD, MGUS and MM LOW. In MM HIGH white arrows point MSC CD271+ IFN+ (GBP2/ISG15)

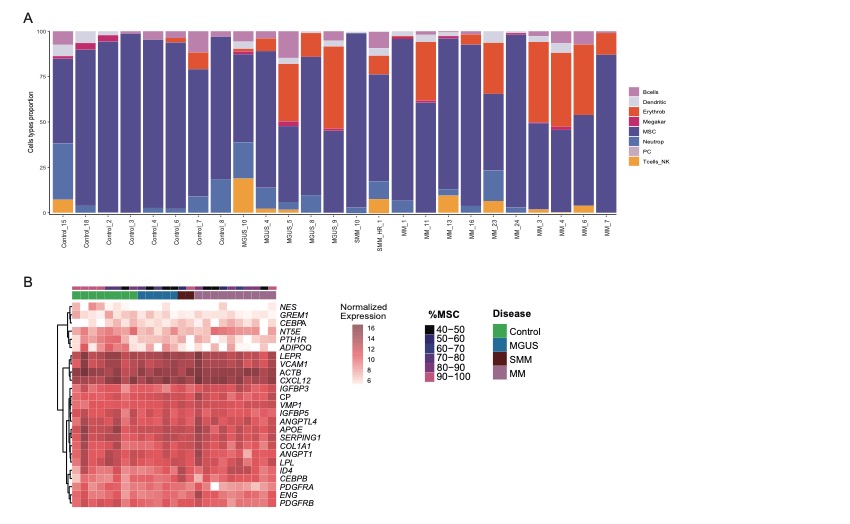
 **Supplemental Figure 15: Evaluation of MSC proportion in bulk RNA-seq human samples.** (A) Bar plot showing the percentage of different cell types present on each sample estimated by CibersortX. (B) Heatmap of MSC canonical markers gene expression correlated with MSC percentage estimated by CibersortX.

SUPPLEMENTAL TABLES

Supplemental Table 1: Description of mice utilized in the study for scRNA-seq and IF analysis.

Supplemental Table 2: Clinical characteristics of human individuals for bulk RNA-seq and IF analysis.

Supplemental Table 3: DEGs between MGUS and control EC.

Supplemental Table 4: DEGs between MM and MGUS EC.

Supplemental Table 5: ORA between all stages in MSC.

Supplemental Table 6: DEGs between MGUS and control MSC.

Supplemental Table 7: DEGs between MM and MGUS MSC.

Supplemental Table 8: Full list of significant L-R pairs involved in EC and PC communication and in MSC and PC communication.

Supplemental Table 9: DEA between VRd-treated and MM EC.

Supplemental Table 10: DEA between treated and MM MSC.

Supplemental Table 11: Comparison between human MGUS-MSC and HD-MSC.

Supplemental Table 12: Comparison between human MM-MSC and MGUS-MSC.

Supplemental Table 13: Comparison between human MM-MSC and HD-MSC.
